# Interpretable computational SHG microscopy enables quantitative tissue phenotyping of cardiac remodeling in nonischemic heart failure

**DOI:** 10.64898/2026.10.21.760498

**Authors:** Reid Master, Wesley Poon, Rania Uzma, Stephan Bannikov, Muthu Kumar Krishnamoorthi, Ashok Veeraraghavan, Hong Zhao, Arvind Bhimaraj, Raksha Raghunathan, Stephen T.C. Wong

## Abstract

Heart failure is characterized by extracellular matrix remodeling and collagen deposition that impair myocardial function. Second harmonic generation (SHG) microscopy provides label-free visualization of fibrillar collagen and myosin but lacks standardized, interpretable, and quantitative frameworks with clinical-level performance. We present a feature extraction pipeline integrating 87 biologically interpretable SHG-derived metrics for automated classification of nonischemic heart failure in murine specimen. The framework distinguished normal, heart failure, and natural recovery states with area-under-the-curve values of 0.84-0.99, outperforming established open-source SHG feature extraction tools. These findings support interpretable, SHG-based quantification of cardiac remodeling and establish a methodology for future multimodal biophotonics applications.

## 1. Introduction

Heart failure (HF) is a clinical syndrome characterized by impaired cardiac pump function and is broadly categorized into ischemic, most commonly due to myocardial infarction, or nonischemic, arising from genetic, valvular, inflammatory, arrhythmic, or hypertensive causes [1]. Clinically, HF presents with fatigue, dyspnea, edema, and organ dysfunction, and remains a leading cause of morbidity and mortality worldwide.

A central feature of HF is cardiac fibrosis, characterized by excessive collagen deposition and crosslinking within the extracellular matrix (ECM) [2]. This process is largely driven by activation of cardiac fibroblasts and disruption of the balance between collagen synthesis and degradation. As collagen accumulates within the interstitial and perivascular regions, the myocardial stiffness increases, impairing both mechanical relaxation and electrical propagation. These structural alterations contribute directly to ventricular remodeling, reduced cardiac output, and progressive decline in cardiac function, particularly within the left ventricle (LV), resulting in systolic and/or diastolic dysfunction [3].

Since fibrosis alters the structural organization of the ECM, the ability to visualize and quantify fibrillar architecture is central to understanding disease progression in HF. Changes in SHG signal density, texture, and spatial heterogeneity may not only influence myocardial mechanics but may also serve as early indicators of pathological remodeling before clinical symptoms develop. These features can provide quantitative markers of tissue state for mechanistic and preclinical studies, although their diagnostic or longitudinal-monitoring value will require validation against histology, cardiac function, and independent cohorts.

Traditional histological approaches such as hematoxylin and eosin (H&E) or Masson’s trichrome staining are widely used to assess fibrotic remodeling and cellular pathology. While these methods reveal critical morphological information including cellular heterogeneity, nuclear characteristics, and other pathognomonic features, they require tissue excision and destructive staining protocols, limiting their potential for real-time or in vivo applications [4]. As a result, there is growing interest in imaging modalities that enable label-free and minimally invasive quantification of collagen structure, particularly those that preserve tissue architecture while enabling quantitative assessment of ECM remodeling.

Second harmonic generation (SHG) microscopy is a label-free nonlinear optical technique that visualizes noncentrosymmetric fibrillar structures, including collagen and myosin without fluorescent dyes or chemical stains [5, 6]. Since cardiac fibrosis is characterized by collagen deposition and remodeling of ECM, SHG microscopy provides a promising biologically nuanced approach for studying structural changes associated with heart failure. Prior studies have demonstrated that SHG can visualize collagen remodeling and detect structural alterations associated with HF in murine models [7-12]. However, these studies have largely focused on qualitative assessment or a limited number of task-specific quantitative metrics rather than comprehensive, interpretable characterization of collagen architecture. Consequently, quantitative SHG analysis of collagen remodeling in heart failure remains underdeveloped, in part due to the strong dependence on mathematical and computational approaches to quantify structural patterns within collagen architecture. This study addresses this gap by developing a broad, biologically interpretable feature extraction pipeline that unifies diverse analytical domains into a single quantitative framework for assessing nonischemic heart failure remodeling.

In other tissue types, SHG has steadily progressed towards in vivo applications for disease assessment [13]. With the advent of deep learning (DL) and increasingly advanced ensemble learning techniques designed for tabular data, the performance of these models is poised to approach clinical standards in the near future [14]. However, DL architectures such as U-Net, a widely used convolutional neural network (CNN) in medical imaging, typically require large datasets and substantial computational resources. This has led to an increasing interest in the use of tabular feature inputs for ensemble-based models. These approaches include decision trees (DT), linear regression models, and related algorithms. The “black box” nature of DL models can also limit biological interpretability, which is critical for clinical decision making, particularly for SHG, which is less familiar to many pathologists and clinicians. Thus, there is a need for a streamlined, high-throughput pipeline capable of extracting interpretable, handcrafted feature sets from individual fields of view (FOVs), while ensuring that model validation is performed at the specimen level rather than treating correlated FOVs as independent biological samples.

Several analytical tools have been developed as precedents, including CurveAlign and CT-FIRE [15] as well as TWOMBLI [16]; however, these approaches typically capture only a limited portion of the variability present in a given dataset and often focus primarily on fiber-level features such as alignment and orientation. Furthermore, their generalizability across diverse tissue types and imaging conditions remains limited. Such feature engineering approaches can be leveraged to extract biologically meaningful information from SHG image stacks to be input into various machine learning (ML) frameworks, including random forest (RF) classifiers [17]. Alternatively, these handcrafted features may also be used for unsupervised FOV classification [18].

Numerous studies have applied quantitative SHG microscopy to extract features that perform well for specific tasks (e.g., sarcomere length quantification [19]), but many such workflows either depend on idealized FOVs or provide a limited subset of structural descriptors. In this work, we present an automated SHG quantification framework to evaluate fibrotic remodeling in a murine nonischemic HF model and a therapy-associated remodeling state after natural recovery (NR). To the best of our knowledge, the proposed framework represents one of the most comprehensive SHG feature extraction approaches for cardiac remodeling to date, integrating analytical methods from prior SHG and texture-analysis studies into a modular, interpretable feature set.

## 2. Methods

### 2.1 Animal Models

Nonischemic HF was induced in 3-month-old C57BL/6 mice via administration of drinking water supplemented with 1% sodium chloride and 0.3 mg/mL L-NG-nitroarginine methyl ester (L-NAME), a nitric oxide synthase inhibitor, combined with continuous infusion of angiotensin II (0.7 mg/kg/day) for 5 weeks using a subcutaneous osmotic pump. Age-matched control mice received no L-NAME or angiotensin II exposure. For the recovery group, HF-inducing agents were withdrawn at week 5, and mice were subsequently maintained on regular drinking water for an additional 4 weeks (weeks 5-9) to allow for NR. All procedures were performed under an approved protocol by the Houston Methodist Research Institute Institutional Animal Care and Use Committee.

### 2.2 Tissue preparation

Following euthanasia, the heart tissue was collected, fixed in formalin, embedded in paraffin, and sectioned into 5 µm slices. One unstained section was imaged using SHG microscopy, while the adjacent serial section was stained with Masson’s trichrome and imaged using bright-field microscopy.

### 2.3 SHG image acquisition

A dual output ultrafast tunable laser (Insight DeepSee, Spectra-Physics, Santa Clara, USA) was used as the light source for the SHG system. Samples were imaged using a Bergamo multiphoton microscope (Thorlabs, Inc. NJ, USA) at 25X magnification with the tunable source of the laser set to 803 nm. An emission filter of 400 ± 10 nm was used to filter the light from the sample, and a photomultiplier tube was used to collect the light. SHG signals were acquired in the backward direction for analyses.

For each tissue specimen, whole-slide SHG images were acquired. Each FOV consisted of a 512 × 512 pixel grid, with a pixel size of 0.94 μm, corresponding to an imaging area of approximately 430 μm × 430 μm per FOV. At each FOV, images were acquired along the entire depth of the slide and at each depth 4 frames were acquired and averaged to improve the signal-to-noise ratio. Finally, a maximum intensity projection (MIP) was subsequently generated across the depth stack for each FOV along depth, and the resulting MIP image was used for quantitative analysis. As a preprocessing control, a region of the glass slide without tissue was imaged after completing data acquisition for each specimen.

### 2.4 Preprocessing

Figure 1 provides an overview of the complete image analysis pipeline, from data acquisition through feature extraction to ML classification. Steps 1-5 in Figure 1 depicts the preprocessing steps. In summary, raw SHG tiles first undergo BaSiC shading correction to compensate for nonuniform illumination, using estimated flatfield and darkfield components [20]. Each corrected tile is subsequently median filtered to reduce salt-and-pepper noise. Background subtraction is then performed using a threshold defined as the 99th percentile intensity of a glass-only FOV, with negative values set to zero. The resulting image stack is globally normalized through histogram stretching and contrast enhancement prior to export of preprocessed tiles and corresponding montages.

**Fig. 1.**
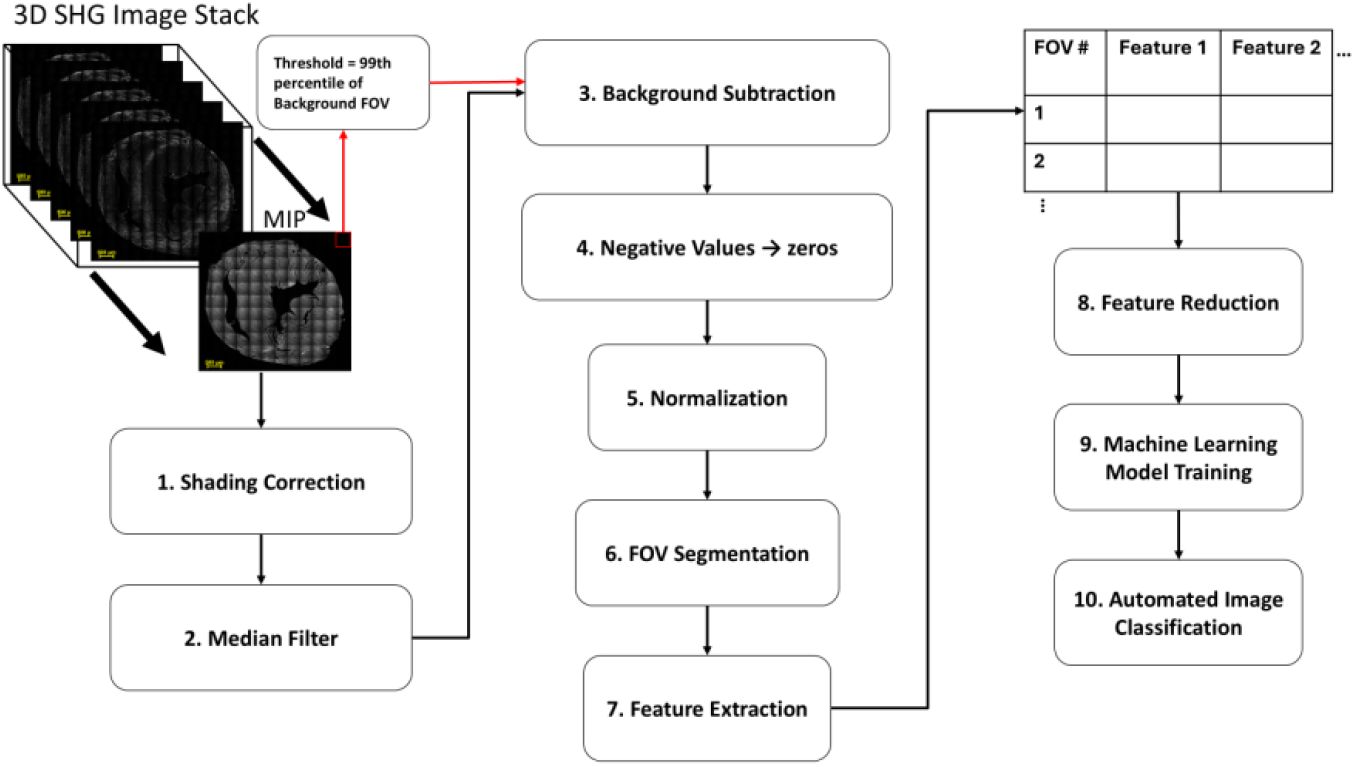
Summary of SHG feature extraction pipeline.

### 2.5 Segmentation

Given that the sample sets were unannotated and the distinction between normal and failing hearts were not defined at the FOV level, a specific anatomical region was segmented to reduce confounding variability and ambiguity in interpretation. The left ventricular (LV) free wall was selected due to its well-established susceptibility to fibrotic remodeling in heart failure [2]. For each heart sample, five FOVs were selected from the LV free wall from the preprocessed SHG montage based on the following criteria.

1. Collagen fibers and myosin filaments were oriented longitudinally to minimize sampling variability arising from differences in axial tissue sectioning.
2. FOVs were directly adjacent to one another.
3. Only FOVs from the LV free wall were included to assess region-specific anatomical variation without introducing selection biases.

This controlled ROI strategy improves consistency for proof-of-concept evaluation but should not be interpreted as random or exhaustive sampling of the entire LV.

Figures 2A-C present representative whole-slide SHG images from the control, HF, and NR groups, respectively, with red squares indicating the selected FOVs for sample (segmentations for all 15 murine samples can be found in the supplementary document, Figures S1-S15). Figure 3 displays the corresponding individual FOVs from one sample (depicted in Figure 3), with the top row representing the control group, the middle row the HF group, and the bottom row the NR group.

**Fig. 2.**
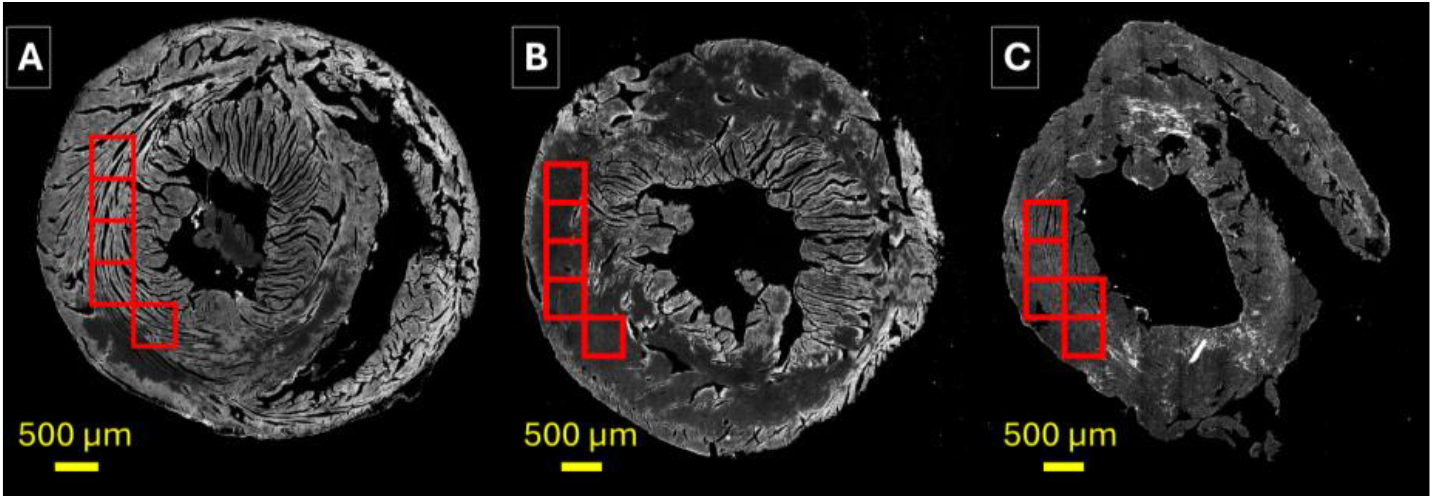
Whole slide SHG images of murine cardiac cross sections from control (A), heart failure (B), and natural recovery (C) models with segmented FOVs.

**Fig. 3.**
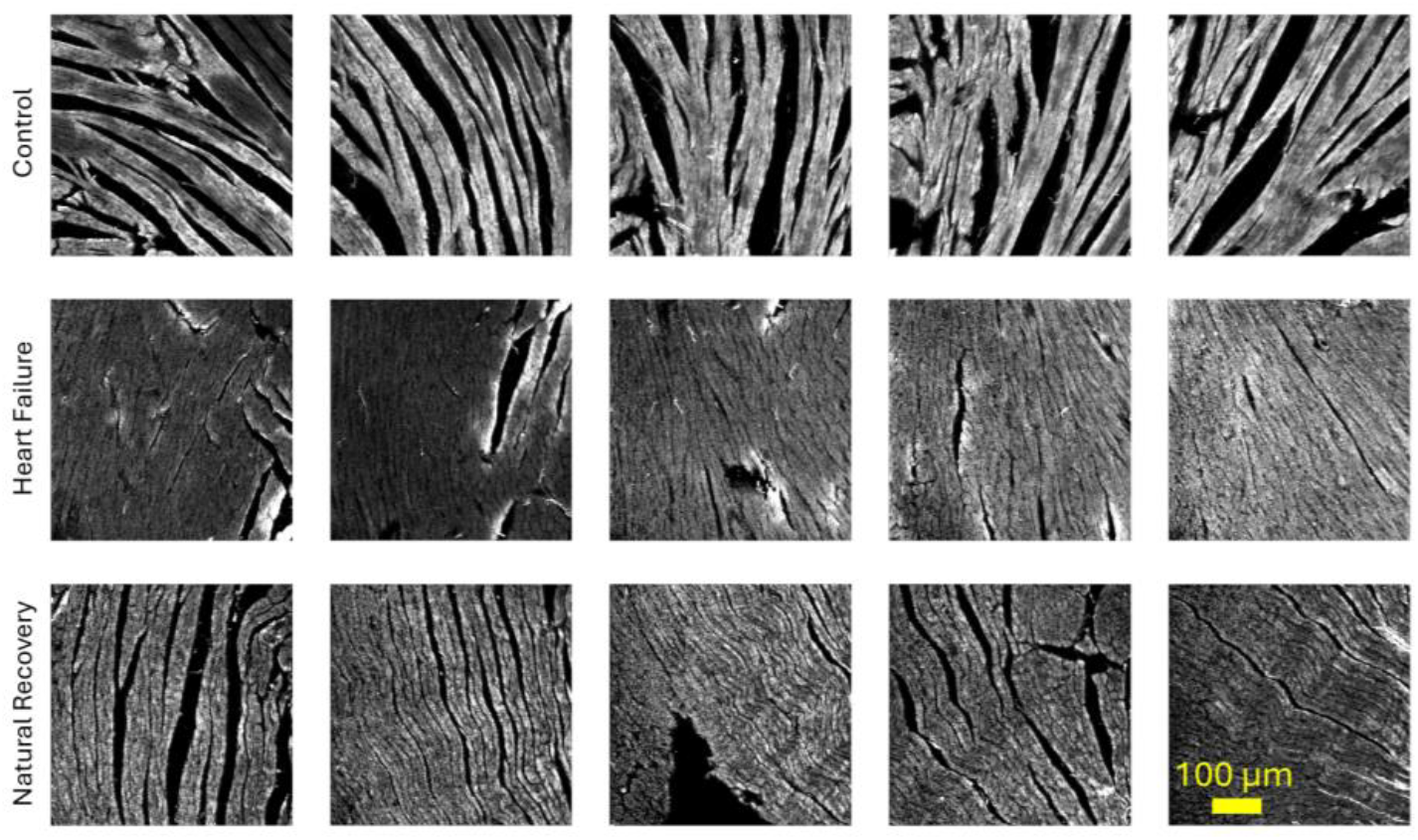
Sample FOVs from control (top), heart failure (middle), and natural recovery (bottom).

### 2.6 Feature extraction

Feature extraction was performed on the selected individual FOVs from each sample. Handcrafted single-metric features were designed to create an interpretable parameter set with high class-separability suitable for input into supervised ML models. To reduce redundancy, only informative and interpretable features from each analysis domain were retained. All selected metrics were constrained to have low computational cost to support potential clinical deployment. To ensure robustness to arbitrary imaging orientation, all parameters were developed and optimized to be rotationally invariant wherever applicable (e.g., textural features) by averaging across four canonical directions, and orientation-dependent metrics (e.g., mean preferred orientation angle) were excluded. This framework supports high-throughput, quantitative SHG analysis while preserving feature-level biological interpretability.

To capture spatial heterogeneity in a quantitative, tabular format, a range of computational analyses were performed, many of which have been leveraged in previous studies [21, 22] and carry interpretable biological meaning [23]. These methods were grouped into four primary categories: pixel-level features, textural analysis, local collagen morphology, and global collagen morphology. The specific approaches are further subdivided and summarized in Table 1, yielding a total of 87 tabular SHG features per FOV.

**Table 1.**
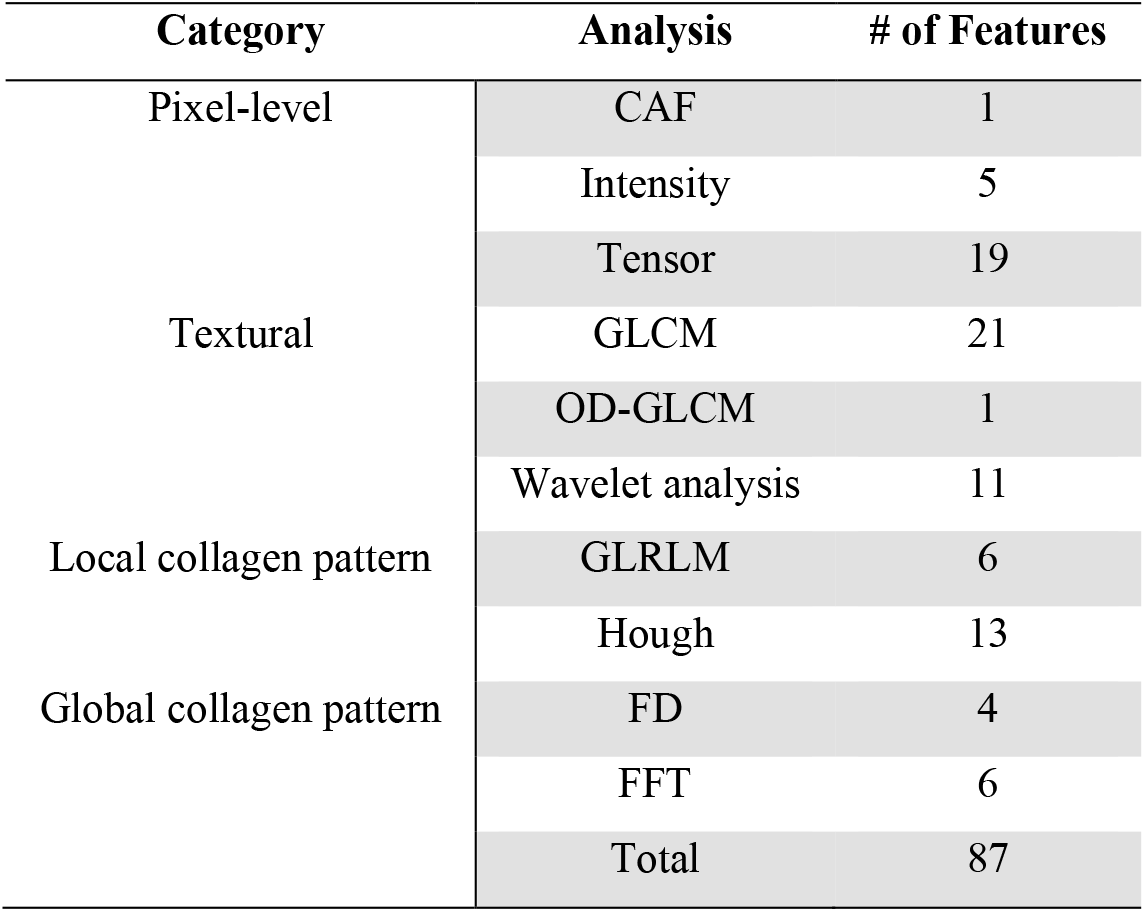
Summary of features by group.

#### 2.6.1 Pixel level analysis

The initial and most fundamental analyses were performed at pixel level, including raw SHG signal intensity-based and tensor-based analysis of this signal. These metrics enable quantitative assessment of tissue patterns directly within the spatial domain.

#### 2.6.2 Intensity analysis

Histogram-based analysis was performed to assess the SHG signal intensity within each FOV. Extracted metrics included mean, standard deviation (std), skewness, kurtosis, and Shannon entropy [18, 24, 25]. Collectively referred to as first-order statistics (FOS), these features provide a quantitative representation of intensity distributions and are used throughout this study to encode histogram-derived information into interpretable tabular parameters. A total of 5 features were derived: *intensity mean, std, skewness, kurtosis*, and *entropy*.

#### 2.6.3 Collagen area fraction

As previously described [22], collagen area fraction (CAF) represents the relative area occupied by collagen within a given FOV. Areas of fibrosis with high rates of collagen deposition tend to have a higher CAF, however, in cardiac tissue, this feature should be interpreted as SHG-positive fibrillar area fraction because the binary signal cannot distinguish collagen from myosin without additional validation or polarization-sensitive acquisition.

#### 2.6.4 Tensor analysis

Structure tensors form the backbone of numerous image processing algorithms and have been widely used for quantitative SHG studies [26-30]. In this approach, each intensity matrix is first converted into gradient estimates using MATLAB gradient, followed by mild Gaussian smoothing. The 2×2 structural tensors are then calculated for each pixel as

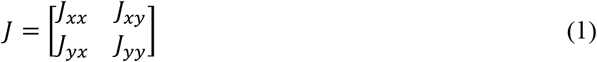

where *J*_*xx*_ = *G*_*σ*_ (*I*_*x*_)^2^, *J*_*yy*_ = *G*_*σ*_ (*I*_*y*_), and *J*_*xy*_ = *J*_*yx*_ = *G* _*σ*_ (*I*_*x*_*I*_*y*_)^2^. Here *I* _*x*_ and *I* _*y*_ denote the horizontal and vertical partial derivatives of pixel intensity *I*, respectively, and *G*_*σ*_ represents Gaussian smoothing with scale parameter. From the fundamental tensor components, corresponding eigenvalues are derived (Eq. 2).

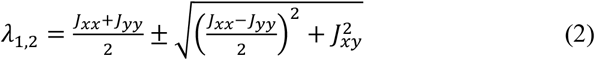

Then, pixelwise degree of isotropy (DOI), tensor gradient energy (trace), and preferred orientation (PO) were calculated (Eqs. 3-5).

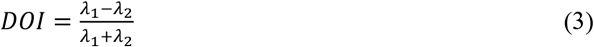

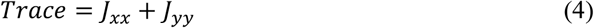

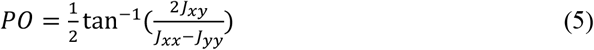

Finally, anisotropy-weighted intensity (AWI) is defined as the matrix multiplication of DOI with pixel intensity. Because gradient-based measures are nondiscriminatory to low-intensity background regions, this metric emphasizes areas exhibiting both strong directional organization and high signal intensity, thereby correlating closely with collagen fiber scaffolding. These tensor-based metrics are calculated for each pixel from each SHG FOV, generating corresponding pixelwise heatmaps. From corresponding histograms, the same FOS features used in intensity-based analyses: mean, std, skewness, kurtosis, and entropy are extracted. The mean PO metric is not reported, as it is inherently dependent on image orientation. Further, orientation is a periodic, circular variable rather than a linear one. Collagen fiber orientation lacks polarity, i.e. fibers aligned at 0° are approximately the same as fibers aligned at 180°. Similarly, fibers aligned at 1° are approximately the same as fibers aligned at 179°. In such situations, a linear metric would incorrectly treat them as maximally different. Hence computing the std using a circular formulation with a double-angle transform accounts for this periodicity by mapping orientations onto the unit circle, ensuring that angular wrap-around is handled correctly. This yields a dispersion measure that reflects true variability in fiber alignment rather than artifacts introduced by arbitrary angular boundaries. Thus, the std of PO is computed using circular, rather than linear formulation (Eq. 6).

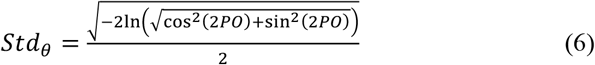

#### Biological interpretability

Tensor-derived features offer strong biological interpretability compared to more abstract textural metrics. For example, mean AWI, reflects the relative abundance of bright, well-aligned fibers while suppressing dim or isotropic regions, and is therefore expected to be elevated in collagen-dense, fibrotic environments. In parallel, FOS descriptors of AWI capture variation, asymmetry, and heterogeneity within these maps. DOI quantifies directionality independent of intensity, providing complementary information in cases where signal intensity is noisy. Additional tensor features, such as trace (reflecting overall gradient energy and structural contrast) and preferred orientation (capturing local angular relationships), further characterize directional organization among neighboring pixels.

Collectively, these metrics quantify both intensity-weighted and directional properties of collagen architecture. When integrated with morphological and textural features, this framework enables comprehensive representation of both local patterns and global structural organization, facilitating the conversion of SHG images into interpretable, quantitative descriptors [21]. A total of 19 features were reported from tensor analysis, including *std, skewness, kurtosis*, and *entropy* for each *AWI, DOI, trace*, and *PO. Mean* was also reported for *AWI, DOI*, and *trace*.

#### 2.6.5 Texture analysis

Texture-based analyses, such as gray-level co-occurrence matrix (GLCM) and wavelet decomposition, have been extensively applied across diverse tissue types and SHG imaging applications [24-26]. Although individual metrics derived from these methods may lack direct biological interpretability, their combined use provides a powerful framework for classifying FOVs [18, 31].

#### 2.6.6 Gray-level co-occurrence matrix (GLCM)

GLCM-based feature extraction yielded a total of 21 features. These include *autocorrelation, contrast, dissimilarity, homogeneity, maximum probability, GLCM energy, GLCM entropy, cluster prominence, cluster shade, cluster tendency, correlation, sum of squares variance, inverse variance, sum distribution average, sum distribution entropy, difference distribution entropy, difference distribution variance, information measures of correlation 1, information measures of correlation 2*, inverse difference normalized (*invdn*), and *inverse difference moment normalized*. Full details of their calculation, including formulae, are described in our previous publication, in which these features were developed and applied to quantify second-order textural characteristics of thyroid neoplasms [22]

GLCM metrics such as contrast, dissimilarity, homogeneity, energy, entropy, and maximum probability characterize intensity variation, smoothness, and uniformity based on gray-level co-occurrence. Correlation and autocorrelation quantify linear dependencies between gray levels, while cluster prominence, shade, and tendency capture joint moments of intensity pairs. Additionally, sum and difference matrices were evaluated through their respective averages, variances, and entropies. Information measures of correlation assess the mutual dependence between marginal distributions, and inverse difference metrics describe local homogeneity with reduced sensitivity to gray-level variation. All features were computed from normalized probability distributions.

#### 2.6.7 Orientation dependent gray-level co-occurrence matrix (OD-GLCM)

OD-GLCM analysis was used to assess rotational variance of GLCM-derived cumulative correlation, thereby characterizing global directionality. This analysis contributed one metric, *std of cumulative correlation* [22], which was calculated to approximate textural anisotropy. Higher values indicate directionality which varies with orientation while lower values suggest isotropic behavior which is rotationally invariant. Complete details of this analysis are described in our previous work.

#### 2.6.8 Wavelet transform

Discrete wavelet transform analysis was performed to quantify the multiscale organization and frequency dependent complexity of collagen structures in SHG images [32-34]. By capturing both spatial and frequency-domain information, wavelet-based methods enable assessment of hierarchical collagen architecture that is not readily resolved using purely spatial metrics.

Each grayscale SHG FOV underwent a two level 2D wavelet decomposition using the MATLAB functions *wavedec2* and *detcoef2*, with a built in MATLAB wavelet such as *Daubechies db4*. The decomposition separated the image into one approximation sub band (LL), capturing low frequency structural content, and three detail sub-bands (LH, HL, HH) representing horizontal, vertical, and diagonal high frequency information, respectively, at each level. Level 1 captures fine scale fibrillar structures, including collagen fibers and edges, while level 2 captures coarser architectural patterns such as fiber bundles or perivascular networks. From this multiresolution framework, we computed a set of global scalar descriptors reflecting energy distribution and textural entropy. In addition to *approximation mean, energy*, and *entropy*, 8 additional metrics were calculated: L1 detail energy (*L*1_*total*_), L2 detail energy (*L*2_*total*_), total detail energy (*E*_*total*_), fine to coarse energy ratio (*Ratio*_*L*1/*L*2_), approximation to detail energy ratio (*Ratio*_*LL*/*details*_), L1 detail entropy (*Ent*_*L*1_), L2 detail entropy (*Ent*_*L*2_), and global *wavelet entropy* (Eqs. 7-14). Here *e* denotes sub band energies.

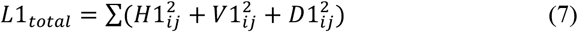

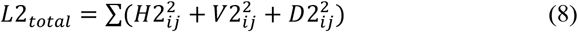

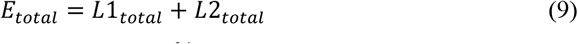

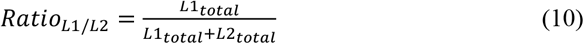

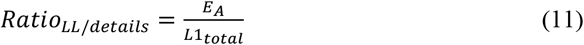

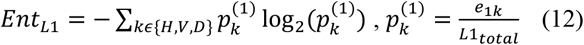

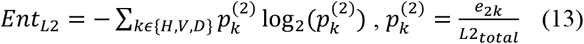

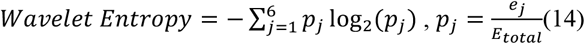

Wavelet derived metrics collectively quantify how energy and complexity are distributed across spatial scales and orientations. A high L1 fraction, low L1 entropy, and/or low L2 entropy indicates organized, fine scale anisotropy typical of aligned collagen fibers, whereas elevated wavelet or approximation entropy reflects isotropic or disordered matrices characteristic of fibrotic remodeling. The approximation to detail ratio distinguishes smooth background regions from highly textured fibrillar domains. Together, these parameters summarize hierarchical collagen organization within each SHG image, complementing pixel-based intensity, tensor, and GLCM analyses to yield a comprehensive multiscale textural profile of extracellular remodeling. Thus, we derive a total of 11 handcrafted, tabular features from wavelet analysis.

#### 2.6.9 Local collagen morphology

To complement these analyses, we evaluated local collagen morphology by quantifying both spatial heterogeneity and larger-scale structural patterns. Several open-source toolkits operate in this realm; notably, TWOMBLI has been used to derive metrics such as fractal dimension and lacunarity (lac) [16], while FiberFit quantifies radial anisotropy using Fourier transform-based approaches. Although these tools demonstrate high-throughput capabilities, they are limited to relatively narrow feature sets [35].

#### 2.6.10 Gray-level run length matrix (GLRLM)

GLRLM analysis has been previously applied in SHG to quantify collagen organization at both local and global scales by measuring the occurrence of contiguous pixel runs within specified gray-level ranges across defined directions (0°, 45°, 90°, and 135°) [18]. As with GLCM and OD-GLCM, we ensure rotational independence of FOV inputs by taking the average of the four canonical directions. Using the built in MATLAB function, *grayrlmatrix*, 11 features were extracted: short run emphasis (*SRE*), long run emphasis (*LRE*), gray-level nonuniformity (*GLN*), run length nonuniformity (*RLN*), run percentage (*RP*), low gray-level run emphasis (*LGRE*), high gray-level run emphasis (*HGRE*), short run low gray-level emphasis (*SRGLE*), short run high gray-level emphasis (*SRHGE*), long run low gray-level emphasis (*LRLGE*), and long run high gray-level emphasis (*LRHGE*). A run is a maximal sequence of pixels which are both contiguous and collinear with identical gray level *i* and run length *j*. The GLRLM function *P*(*i,j*) simply counts the number of occurrences at gray level *i* and run length *j*. The probability normalization function is defined (Eq. 15) and used for subsequent calculations.

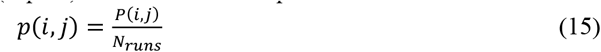

Directional GLRLMs were computed independently and subsequently averaged. Of the aforementioned 11 features, 6 were selected for further analysis based on their high interpretability and prognostic value: LRE, GLN, RLN, RP, HGRE, and LRHGE (Eqs. 16-21).

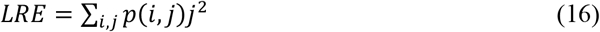

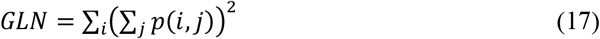

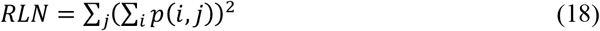

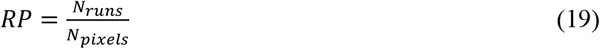

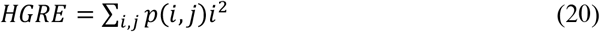

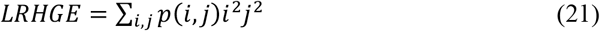

#### Biological interpretability

GLN and RLN capture variability in gray-level distribution and run length structure, corresponding to tissue heterogeneity. LRE, HGRE, and LRHGE each emphasize patterns with high intensity and/or long runs, measuring continuity of collagen fibers in SHG images.

Metrics which emphasized short runs and low gray level occurrences (SRE, LGRE, SRGLE, SRHGE, and LRLGE) were excluded from morphological analysis given that such calculations would primarily detect background and/or random noise with little biological interpretability or significance, though such calculations could still prove useful in textural analysis. Hence, a total of 6 features were calculated using GLRLM analysis.

#### 2.6.10 Hough transform

Building on our prior work [22], we extended Hough transform-based analysis to quantify global collagen alignment by constructing a spatially resolved orientation field and associated coherence map. The goal of this approach was to estimate both the dominant fiber orientation (Hough angle) and the reliability of that orientation (coherence) within localized regions of each SHG image.

Each FOV was first edge-enhanced using Canny detection and partitioned into overlapping square windows (51 × 51 pixels, step size 32 pixels). In prior work, we used 10-pixel steps; however, because this feature had substantially longer run times than other features, we used a 32-pixel step in the present study to balance spatial resolution with computational efficiency. For each window, Hough transform was computed to obtain an angular distribution of line orientations. The dominant orientation, or Hough angle (θ), was defined as the angle corresponding to the peak of the smoothed Hough accumulator. As in our prior work, we once again reported Hough std as a single feature by averaging between all 51 × 51 pixel windows within a given FOV [36].

From this local analysis, we construct a line field (Figure 4A), where each valid window is represented by a short line segment centered at the window location and oriented at the Hough angle, θ. Because the Hough transform returns the orientation of the line normal rather than the line itself, 90 degrees was added to recover the true fiber axis, and the resulting angle was wrapped to the range [0, π) to enforce axial symmetry. A coherence value is also assigned to each window based on the spread of its Hough angular distribution. Specifically, coherence is defined as an inverse function of the windowed Hough std, such that windows with a narrow, sharply peaked angular spectrum yield high coherence (strong alignment), while those with broader spectra yield low coherence (disorganized or uncertain alignment).

**Fig. 4.**
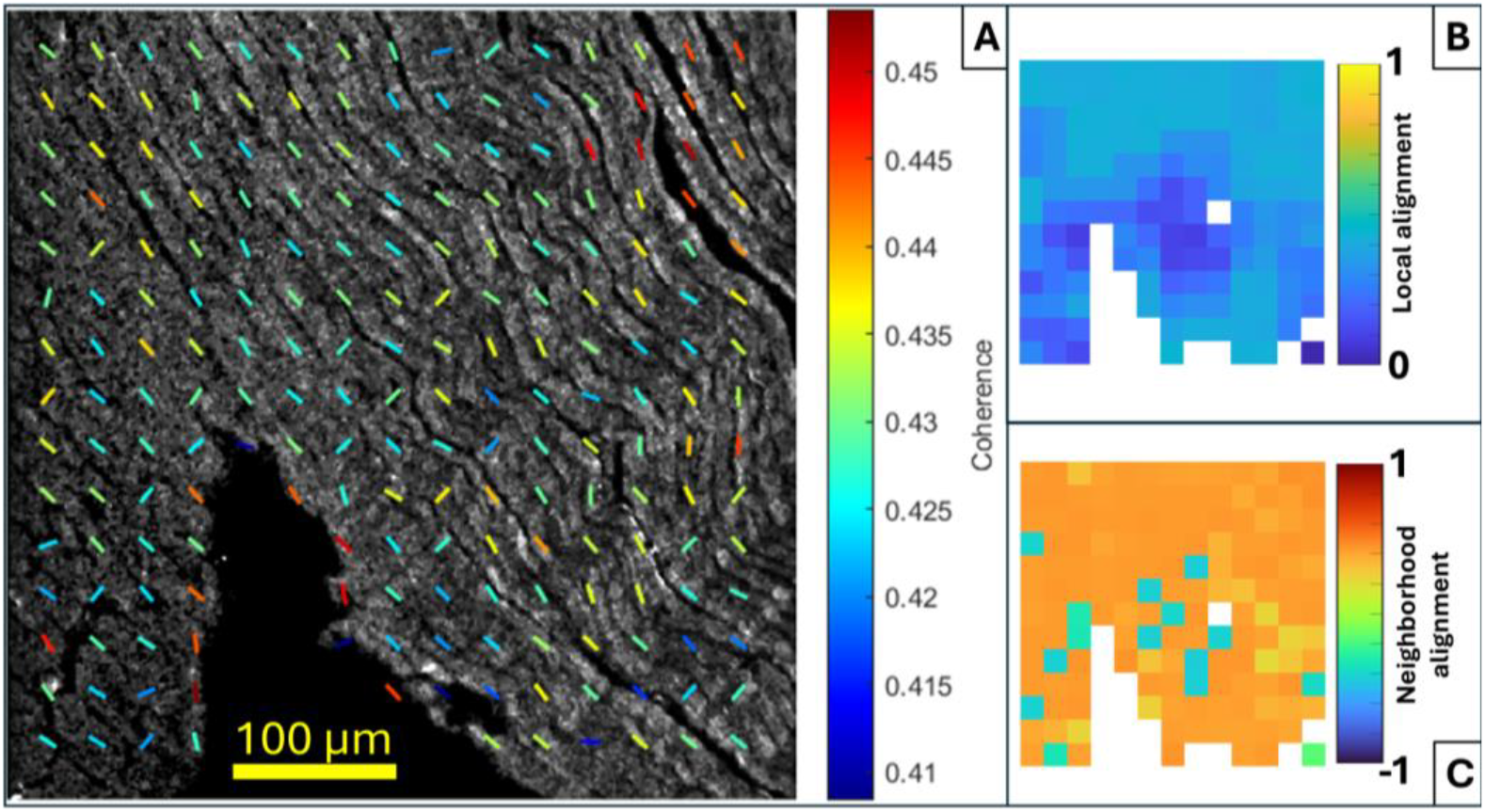
Sample Hough field map (A) using 51×51 pixel windows at a 32-pixel step. Lines are overlaid on the SHG image and oriented at the local Hough angle. Colors reflect each window’s associated coherence value. Corresponding heat maps for local alignment (B) and neighborhood alignment (C).

Figure 4A illustrates this concept: each colored line represents the dominant orientation within a 51 × 51 window, while the color encodes coherence.

As shown in Figure 4A, windows with higher coherence are marked by fibers with high intensity, directionality, and/or strong edges. In contrast, windows with lower coherence tend to visually show less directionality, which in our cardiac samples may reflect axial rather than longitudinal tissue orientation. Windows with insufficient edge content were excluded based on a predefined threshold established from prior SHG studies [22]. In the case of our cardiac samples, collagen and myosin fibers tend to be at least somewhat aligned due to one of our selection criteria being longitudinally oriented cross sections.

Additional exclusion criteria were then applied prior to feature extraction. Specifically, a neighborhood consistency mask required each window to have a minimum of eight valid neighboring windows, thereby eliminating isolated or poorly supported measurements. Further, a coherence-based adaptive threshold was applied to automatically remove windows with weak or unreliable orientation estimates. Together, these steps ensure that subsequent metrics are computed only from spatially consistent and structurally meaningful regions, reducing the influence of noise and improving the stability of local alignment measurements.

*Coherence fraction*, defined as the proportion of valid windows exhibiting nonzero coherence after masking, quantifies the fraction of the analyzed FOV containing windows with measurable and reliable orientation estimate. This is computed as the number of valid windows with nonzero coherence divided by the total number of valid windows within the FOV after all masking and exclusion criteria are applied. Thus, coherence fraction reflects the spatial extent of structurally organized collagen and/or myosin relative to the total analyzable region, rather than a direct pixel-level area measurement. Using the dominant Hough angle θ and corresponding coherence value for each valid window, each orientation θ was mapped to (sin(2*θ*), cos(2*θ*)) and multiplied by its coherence to derive a weighted axial vector field. This ensures that regions with strong fiber alignment contribute more significantly to the final feature set while also treating orientations separated by 180 degrees equivalently.

The coherence-weighted resultant vector was then computed for each window *i* (Eq. 22) with orientation *θ* and coherence weight *w*. Orientation dispersion (in degrees) was then calculated from the resultant length *R* using an axial circular std (Eq. 23), where larger values indicate greater directional heterogeneity and smaller values indicate tighter global alignment.

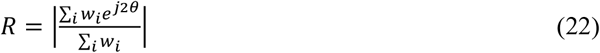

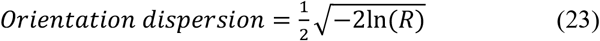

Local organization was quantified by using a 3 × 3 neighborhood averaging scheme. For each central window, represented by a single line in Figure 4A with known dominant angle and coherence value, the doubled-angle vectors from all nine valid windows within the neighborhood, were averaged using coherence as weights, producing a single representative local axial orientation parameter *S*_*local*_ (Eq. 24).

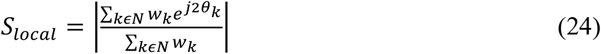

Here, *θ*_*k*_ and *w*_*k*_ denote the Hough angle and coherence weight, respectively, of neighbor *k*, and *N* denotes the 3 × 3 neighborhood. *S*_*local*_ ranges from 0 (locally disordered or multidirectional fibers) to 1 (strong local alignment). Thus, each 3 × 3 neighborhood yields one scalar value summarizing the consistency of fiber orientations within that region. Note that windows along the perimeter of the FOV are handled by our exclusion criteria requiring eight valid neighboring windows; thus, a 51 × 51 pixel window is cropped from the perimeter to reduce inaccurate calculations due to insufficient neighbors. As shown in Figure 4B, regions with consistent fiber orientation exhibit higher *S*_*local*_ compared to regions with less orderly orientations. Finally, neighbor alignment was computed as the agreement between each window’s orientation θ and its local neighborhood mean orientation *θ*_*local*_ defined as the average orientation across all 9 local windows (Eq. 25).

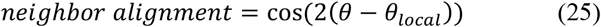

This metric ranges from -1 (opposing orientations) to 1 (perfect alignment). As illustrated in Figure 4C, high neighbor alignment indicates strong agreement with the local structural context, while lower values reflect deviations from local organization. Together with the prior global Hough std metric, these features provide complementary global and spatially resolved measures of collagen directionality, coherence, and regional alignment heterogeneity.

In summary, 13 features were reported using this analysis including *Hough std, coherence fraction, orientation dispersion in degrees*, as well as *mean, std, skewness, kurtosis*, and *entropy* of each *S*_*local*_ and *neighborhood alignment*.

#### 2.6.11 Global collagen morphology

Global collagen morphology characterizes whole FOVs based on large scale patterns which are best captured by fractal dimension (FD) and Fourier analyses. This category of features differs substantially between tissue types and is influenced by factors such as collagen density, recurrent structural motifs, and global collagen alignment.

#### 2.6.12 Fractal dimension

To quantify collagen complexity and spatial motifs, we implemented the box counting method to extract FD [24, 37, 38] and lac [25]. FD can measure self-similarity of biological structures across an exponentially varying scale. Higher FD indicates more irregular, space-filling collagen architecture and the persistence of fine structural detail across progressively smaller spatial scales. Generally, lower FD implies simple, smooth, or less complex architecture overall. Lac, on the other hand, maps tissue heterogeneity via gap distributions across differing scales. Thus, low lac suggests uniform, evenly filled texture while high lac represents patchiness or irregular voids within the collagen network. Both binary and grayscale formulations were evaluated for each metric to assess both structural and textural patterns.

Images were first tiled into nonoverlapping *i* × *i* dimension boxes where box size *i* ϵ[2^0^, 2^1^, … 2^*n*^]. Box sizes start at a single pixel and continue to double in size until the entire 512 × 512 image is occupied. FD (Eq. 26) and lac (Eq. 27) were derived from moments M1 (Eq. 28) and M2 (Eq. 29). Here, *Q* is the total number of *i* × *i* boxes tiling the image, *X* is the number of nonzero pixels in box *k* (for binary calculations) and local std of intensities in box *k* (for grayscale calculations). The first moment depicts the average occupancy (binary) or std (grayscale) content per box, and the second moment takes the squared average to emphasize larger occupancy or variation. For binary FD calculations, boxes containing a nonzero pixel after binarization are recorded as *N(i)*, while the grayscale FD calculation denotes *N(i)* as having a nonzero std.

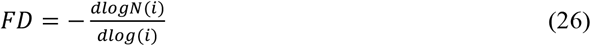

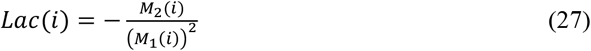

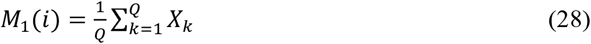

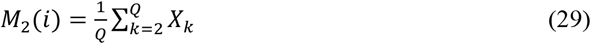

*Binary* and *grayscale* calculations for each *FD* and *lac* yielded a total of 4 features.

#### 2.6.13 Fourier analysis

Most studies that have used FFT-derived features for SHG analysis primarily focus on identifying the dominant fiber orientation within an FOV or quantifying the associated coherence [37, 39-41]. These features have demonstrated statistical significance across a range of tissues and disease states. However, metrics such as aspect ratio (AR) are generally most informative in highly specific contexts where fibers are densely packed and strongly aligned [23]. To reduce reliance on idealized FOVs and enhance generalizability, our Fourier analysis focuses primarily on power spectral density-based metrics described in our previous work [22]. Features included *low, middle, high*, and *total frequency power*.

Weighted frequency center (*WFC*) and *anisotropy* are two FFT-based quantitative metrics used for global morphological analysis. WFC is used to measure the dominant frequency present in an FOV, the inverse of which represents the dominant wavelength. Larger WFC values correspond to finer structures and smaller values to coarser structures [21]. Anisotropy was also developed to enable higher generalizability of the AR [42]. Briefly, AR is calculated by applying a threshold to a 2D FFT magnitude spectrum, finding an ellipse of best fit, and reporting the ratio between its length and width. An equal ratio gives a value close to 1, indicating low directionality or anisotropy, while a perfectly aligned, evenly spaced bundle of fibers would be expected to approximate a cohesive line in Fourier space, meaning a value close to zero. Our anisotropy metric leverages similar relationships but instead relies on radial summation from the 2D power spectrum, thereby extending its use beyond a specific subset of FOVs. Larger anisotropic values represent higher global directionality at the dominant angle.

In this study, the anisotropy parameter is computed using the same formulation as our prior thyroid work [22], based on radial line integrals of the Fourier magnitude spectrum followed by vector summation of the normalized angular distribution. Specifically, the FFT magnitude is first processed using periodic plus smooth decomposition to reduce edge artifacts, followed by a nonlinear magnitude scaling. Line integrals are then computed across 0 to π, normalized, and mapped using a doubled-angle representation to enforce axial symmetry. The final anisotropy value is defined as the magnitude of the resultant vector, yielding a bounded metric between 0, or completely isotropic structure, and 1, or anisotropic, highly aligned structure. Importantly, both WFC and anisotropy are computed at the full FOV level and therefore represent global descriptors of collagen morphology. While both global and windowed anisotropy were explored in our prior work, only the global formulation is included here, as it demonstrated improved generalizability in small datasets and reduced sensitivity to local noise and segmentation variability. Therefore, windowed anisotropy mapping was not directly incorporated in this study in the same way. Lastly, this Fourier-domain anisotropy parameter is distinct from pixel-level anisotropy measures derived from tensor analysis, as it captures global directional organization in the frequency domain rather than local spatial gradients.

Thus, a total of 6 features (*low, mid, high, and total frequency power*, as well as *WFC* and *anisotropy*) were calculated and reported in our Fourier analysis. A comprehensive tabular summary of all 87 features can be found in the supplementary materials (Table S1).

### 2.7 Validation

To benchmark our feature extraction algorithm against existing high throughput frameworks, we used CT-FIRE [15], a well-established open-source algorithm, to extract collagen features characterizing individual fibers. Using a curvelet-based methodology similar to ridge detection, the CT-FIRE algorithm segments individual collagen fibers to generate a vectorized map from which metrics from each fiber can be extracted. FOS metrics are then output using histograms for each fiber metric: length, width, angle, and curvature. CurveAlign [15] is a related algorithm that extracts local and global morphological features from SHG images; however, it was not utilized due to interference of myosin fibers with the collagen scaffolding. A total of 23 features were extracted directly from the CT-FIRE output and a detailed breakdown of these features can be found in the supplementary materials (Table S2). Additionally, the open-source ImageJ toolkit TWOMBLI [16] was used to benchmark our algorithm. The software functions similar to ours but is hindered by a small subset of just 8 features including collagen area, length, and curvature, as well as lac, FD, endpoints, branchpoints, and high gray-level uniformity. Outputs of the two open-source feature extraction tools for individual FOVs were tabularized and fed into different post processing ML frameworks (refer to sections 2.8 and 2.9) in identical fashion to our pipeline to assess efficacy of our feature set.

### 2.8 Feature selection

To reduce redundancy among highly collinear metrics, we implemented a correlation-based feature filter that operates within each training fold only. After standardizing all numeric features using z scores, we first ranked features with MATLAB ReliefF on the training data, which produces an importance weight for each metric using a greedy algorithm. Features were sorted in descending order of ReliefF weight, and the correlation filter was then applied in this ranked order so that more informative features were preferentially retained.

A pairwise Pearson correlation matrix was generated to compare all feature pairs [18]. In each case where a pair had a Pearson correlation coefficient (PCC) of greater than 0.9, the feature with a lower ReliefF ranking was eliminated, and the higher-ranking feature was retained. This reduced overall redundancy of features while giving preference to higher quality features which explained a greater degree of variance within each FOV. The resulting subset of nonredundant, high importance metrics formed the input feature set for model training in that fold, and the same feature indices were applied to the held-out test data to avoid data leakage.

Principal component analysis (PCA) was used in parallel as a distinctly separate post processing method to reduce dimensionality and stabilize model training by projecting the filtered features into an orthogonal space. This feature reduction approach was compared to the PCC reduction method and the results were reported. After median imputation and z score standardization within each fold, PCA was fitted on the standardized training features to obtain an orthogonal set of components ordered by the amount of variance they explained. Because the data was already standardized, no additional centering was performed during decomposition. The number of retained components was chosen to capture at least 95 percent of the total variance. The training samples were then represented using their principal component scores, and the held-out test samples were projected onto the same loading matrix. PCA was never recalibrated on test data, ensuring that all dimension reduction steps were learned strictly from the training portion of each fold and preventing information leakage.

### 2.9 Machine learning classifiers

For supervised classification of control, HF, and NR regions, we trained machine learning classifiers using leave-one-sample-out (LOSO) cross validation (CV). For each training fold, one murine cardiac sample was left out to avoid biological fingerprinting, which tends to falsely elevate model accuracy in validation. Within each training set, the feature extraction pipeline described above, including per-fold ReliefF ranking and PCC filtering (Section 2.8), was applied before classifier training. Each test set comprised 5 FOVs from the held-out specimen, corresponding to a single class. Class predictions were generated for individual held-out FOVs, while the independence of the validation design was defined at the specimen level.

Although classes were equally balanced at 5 samples each for control, HF, and NR, LOSO CV created a slight class imbalance. To mitigate this, sample weights were assigned so that each class contributed an equal total weight within the training data, effectively equalizing the influence of control, HF, and NR samples. For each held-out specimen, trained models produced class labels and class probability scores for all FOVs. Predictions were aggregated across folds to compute overall accuracy and one-vs-rest receiver operating characteristic (ROC) area under the curve (AUC). ROC curves and confusion matrices for all classifiers are included in the supplementary material (Figures S16-S31).

An RF classifier trained on 300 trees and a minimum leaf size of 1, sampling approximately the square root of the available features at each split. Individual feature importances were computed using the mean decrease in node impurity. For each split node *t* where feature *j* was used, the contribution to importance is defined as the weighted reduction in impurity (Eq. 30).

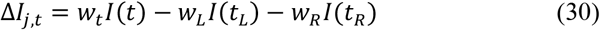

Here, I(*t)* is the node impurity (Gini impurity for classification), *t*_*L*_ and *t*_*R*_ are the left and right child nodes, and *w*_*t*_, *w*_*L*_, *and w*_*R*_ are the proportions of samples reaching each node. The total importance for feature *j* is then obtained by summing Δ*I*_*j,t*_ over all splits involving feature *j* across all trees and averaging across the ensemble to reduce variance (Eq. 31).

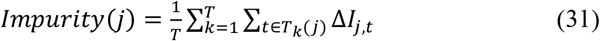

T is the number of trees and *T*_*k*_ (*j*) denotes the set of nodes in tree *k* where feature *j* is used. Raw importance scores were normalized by the sum over all features to yield relative importance values summing to 100%. For interpretability, features were further grouped into predefined analytical domains (e.g., GLCM, wavelet, tensor, Fourier), and group-level importance was computed as the sum of relative importances within each category.

Additionally, we trained multiclass DT, k-nearest neighbors (KNN), and linear discriminant analysis (LDA) classifiers using the same preprocessed feature space, LOSO folds, and class-weighting strategy. The DT classifier (*fitctree*, MATLAB) was fit after feature selection, with tree structure (split variables, thresholds, and depth) determined automatically using the default impurity measure and stopping criteria. The KNN classifier (fitcknn, MATLAB) used k = 7 neighbors, Euclidean distance, inverse distance weighting, and no additional standardization because standardization was already performed during preprocessing. The LDA classifier (*fitcdiscr*, MATLAB) used the *pseudoLinear* option to provide a regularized estimate of the class covariance structure. For each held-out specimen, all classifiers produced class labels and probability scores (posterior probabilities for LDA) for all FOVs, which were aggregated across folds to compute the same performance metrics as the RF model.

Exploratory statistical comparisons of representative features were used to support biological interpretation. Since FOV-level measurements are nested within individual specimens, inferential feature comparisons should be interpreted cautiously unless calculated from specimen-level summaries or modeled with specimen as a random effect. Future validation cohorts should include multiple-comparison correction, confidence intervals, and orthogonal histologic or functional correlations.

## 3. Results

### 3.1 Model performance

Using DT, RF, KNN, and LDA classifiers demonstrated the versatility of our curated tabular feature set. Each model operates on a different ML structure, yet each performed satisfactorily with AUC and raw accuracies for three-class prediction models. Performance of models ranged from 0.84-0.0.99 and 0.76-0.93 for AUCs and raw accuracies, respectively (Table 2).

**Table 2.**
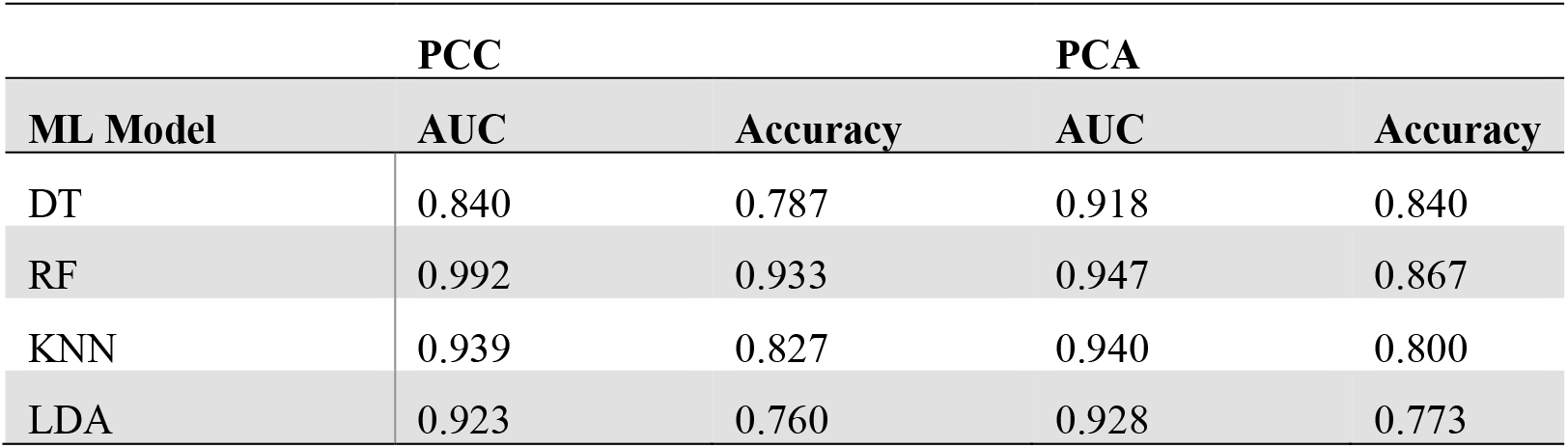
Comparison between machine learning models using feature extraction pipeline.

With respect to our two feature filtering pathways, results demonstrate model-dependent effects of PCC filtering versus PCA. For DT, PCA improved both AUC (0.918 vs 0.840) and accuracy (0.840 vs 0.787), indicating that orthogonalization and variance concentration enhanced split efficiency. In contrast, RF performed best with PCC filtering, achieving the highest overall AUC (0.992) and accuracy (0.933), while PCA modestly reduced performance with an AUC and accuracy of 0.947 and 0.867, respectively. This suggests that ensemble-based tree methods may benefit more from targeted redundancy removal than from rotated feature spaces that obscure original axes. PCC also has the added benefit of direct transparency into importance of individual features in model performance.

For KNN, performance was nearly identical between methods with AUCs and accuracies of 0.939 and 0.827 for PCC vs. 0.940 and 0.800 for PCA, respectively, indicating limited dependency on the specific feature reduction strategy in this dataset. LDA showed slight improvements in AUC (0.928 vs 0.923) and accuracy (0.773 vs 0.760) with PCA compared to PCC, consistent with its reliance on stable covariance estimation and reduced multicollinearity.

Overall, PCC-based ReliefF filtering produced the strongest performance in RF and maintained competitive results in KNN and LDA, while PCA provided measurable gains in DT and modest improvement in LDA. These findings reinforce that optimal preprocessing depends on classifier architecture: tree ensembles may benefit most from correlation pruning that preserves original feature structure, whereas covariance-based linear models and certain axis splitting trees can improve when dimensionality is reduced into an orthogonal, variance-ordered space.

### 3.2 Benchmarking feature extraction algorithm

Table 3 compares performances, summarized by AUC, of different ML models between feature extraction tools using PCC filtering alone for simplicity. ML models trained on our feature set had higher AUCs than those trained on CT-FIRE and TWOMBLI feature sets, with AUCs of 0.84-0.99 compared to 0.34-0.45 and 0.82-0.87, respectively.

**Table 3.**
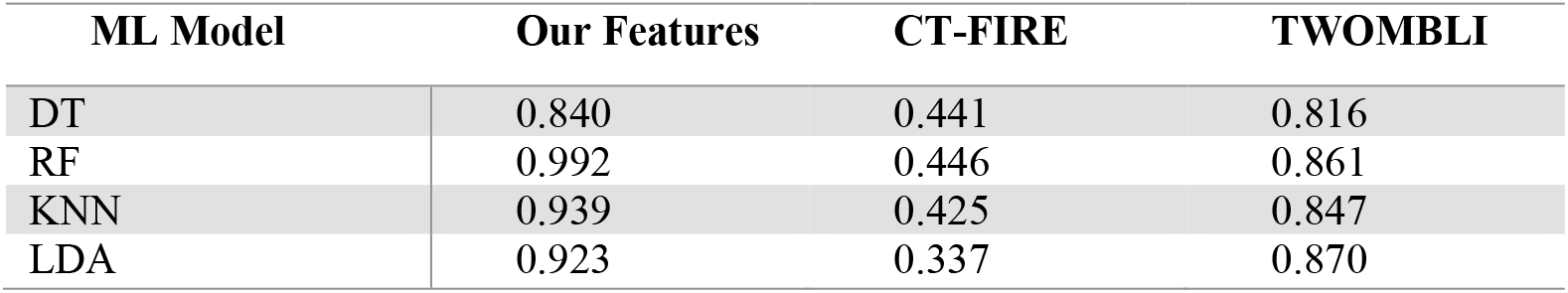
Performance comparison of our feature extraction algorithm with other open-source feature extraction tools.

| ML Model | Our Features | CT-FIRE | TWOMBLI |
| --- | --- | --- | --- |
| DT | 0.840 | 0.441 | 0.816 |
| RF | 0.992 | 0.446 | 0.861 |
| KNN | 0.939 | 0.425 | 0.847 |
| LDA | 0.923 | 0.337 | 0.870 |

### 3.3 Feature importance

To maximize model transparency and biological explainability, we analyzed feature importance using the RF model. Across the ensemble, the model records how much each feature reduces impurity or increases predictive accuracy when used for a split. Averaging these contributions over hundreds of trees yields stable, low variance importance estimates that directly reflect how often and how effectively each feature influences the model’s decisions. Thus, RF is the ideal choice for quantifying feature importance rankings.

Of the 87 initial SHG features, 67 were retained in at least one of the 15 cross validation folds and therefore contributed to the global importance model. The remaining 20 features were consistently removed by the PCC threshold (r ≥ 0.90) due to high redundancy with higher ranked predictors and were excluded from the final importance analysis.

Figure 5A illustrates total importance values based on the type of feature extraction performed (pixel level, textural, local collagen, or global collagen) and is also broken down by the specific type of analysis used in Figure 5B. Textural features dominated total feature importance (55.1%), followed by pixel-level (23.0%), global (12.5%), and local collagen morphology (9.5%). Specifically, these findings were most attributable to features derived from GLCM (32.4%), wavelet (22.6%), and tensor (20.7%) analyses.

**Fig. 5.**
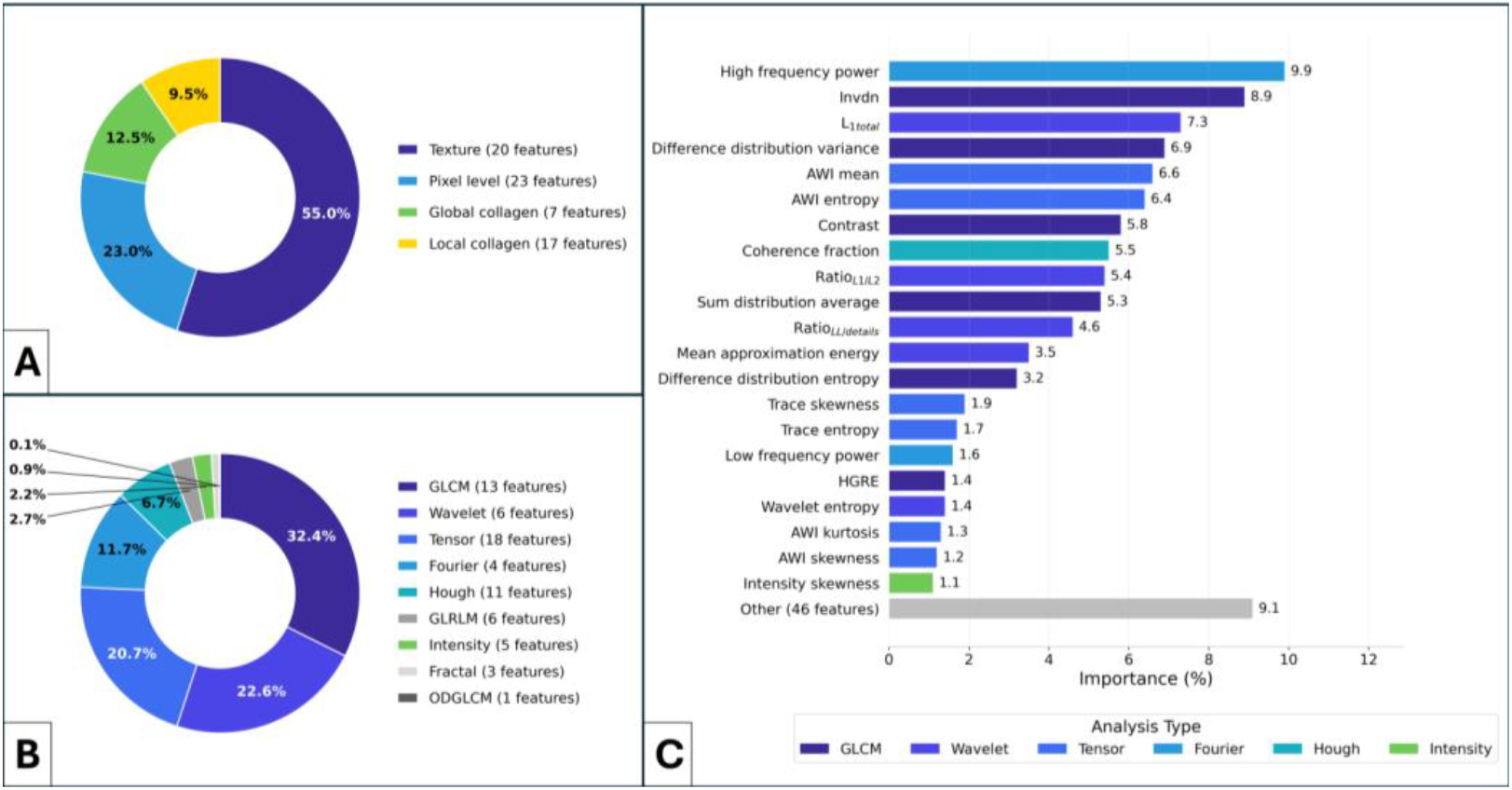
RF importances by analysis group (A), analysis type (B), and individual feature (C).

Figure 5C shows the relative importance of each individual feature among the subset retained after ReliefF ranking and PCC filtering within the LOSO CV framework. High frequency power, a Fourier derived metric, was the top performer, comprising 9.9% of total feature importance. The remainder of the top 10 included features from GLCM, wavelet, tensor, and Hough field analyses. The top 25 also included intensity, GLRLM, and fractal dimension features, demonstrating that discriminatory information is distributed across multiple analytical domains. Importantly, no single feature class dominated performance, and importance was broadly distributed across retained features, underscoring the value of engineering diverse, minimally redundant descriptors to characterize SHG image variance.

PCA can also be broken down into principal component loadings as a measure of feature importance. Although less interpretable, this still adds transparency to models like LDA that natively benefit from PCA dimensionality reduction. A PCA loadings table generated by the same RF model is provided in the supplementary materials (Figure S32).

### 3.4 Biological interpretability

While the feature extraction algorithm presented in this work is generalizable, our primary objective was to assess fibrotic remodeling during heart failure and NR. As stated previously, each metric used in this study is designed to have a clear and interpretable biological meaning. Rather than explaining all 87 features in detail, one representative feature was used from each of the top 5 analysis groups: GLCM, wavelet, tensor, Fourier, and Hough field. The top features for these groups were invdn, L1 total, mean AWI, high frequency power, and coherence fraction, respectively. The performance of these metrics have been tabulated in Table 4. Median values for each murine group can be found in the supplementary materials (Table S3).

**Table 4.** Summary of statistically significant differences between control, heart failure, and natural recovery murine models.

| Feature | Control vs. heart failure | Control vs. natural recovery | Heart failure vs. natural recovery |
| --- | --- | --- | --- |
| Invdn | **** | **** | **** |
| L1 total | **** | **** | **** |
| Mean AWI | **** | **** | * |
| High frequency power | **** | **** | **** |
| Coherence fraction | **** | *** | **** |

\*p < 0.05, \*\*p < 0.01, \*\*\*p < 0.001, \*\*\*\*p < 0.0001

The GLCM-derived feature invdn reflects local homogeneity by quantifying the similarity of neighboring pixel intensities. In the context of SHG imaging, this corresponds to regions where collagen signal is spatially consistent. The observed increase in invdn in HF relative to the control group suggests a shift toward more uniform intensity distributions, potentially reflecting dense homogenized fibrotic deposition. Conversely, the reduction in invdn in the NR group suggests partial restoration of microstructural variability, consistent with disruption or remodeling of this dense collagen network.

L1 total from wavelet analysis captures fine-scale, high-frequency structural content within the image, corresponding to fibrillar detail such as individual collagen fibers and sharp intensity transitions. Reduced L1 energy in HF indicates a loss of fine structural detail, consistent with the replacement of organized fibrillar architecture by more diffuse fibrotic tissue. Notably, the marked increase in L1 total in the NR group suggests re-emergence or reorganization of fine-scale collagen features, potentially reflecting therapeutic remodeling of the extracellular matrix.

From tensor calculations, mean AWI provides a direct and highly interpretable measure of aligned, high-intensity structures by weighting pixel intensity with local anisotropy. Biologically, this metric emphasizes regions where collagen fibers are both bright and directionally coherent, corresponding to structurally intact fibrillar scaffolding. The substantial decrease in AWI in HF relative to control reflects disruption of this organized architecture, while the partial recovery in the NR group suggests re-establishment of aligned collagen structures though not to baseline levels. This trend is consistent with the observed decrease in fine structural detail, L1 total, and increased homogeneity, invdn, in HF, with partial reversal in the NR group, collectively supporting a transition from disorganized fibrotic deposition toward restoration of structured collagen architecture.

High frequency power represents the summed contributions of high frequency content within the Fourier domain and acts complimentarily to wavelet analysis. Similar to L1 total, this metric reflects the presence of small-scale structural features with respect to the global collagen pattern. The reduction in HF indicates a loss of fine, periodic structure in HF tissue, while the significant increase in the NR group suggests increased structural complexity and potentially heightened heterogeneity associated with collagen and/or myosin remodeling. This corroborates with the results of the L1 total metrics. Importantly, the magnitude of this increase may also reflect emerging microstructural irregularity rather than a complete return to normal architecture, highlighting the nuanced interpretation required for frequency-based metrics.

Finally, coherence fraction, derived from the Hough field calculation, directly quantifies the proportion of the FOV exhibiting strong directional alignment. This metric is particularly intuitive, as it reflects the spatial extent of well-organized collagen fibers. The marked decrease in coherence fraction in HF indicates widespread loss of directional organization, consistent with disordered fibrotic remodeling. In contrast, the restoration of coherence fraction in the NR group to levels comparable to control suggests recovery of large-scale alignment patterns, even if finer structural features remain altered.

Taken together, these representative metrics illustrate how distinct analytical domains map onto specific biological characteristics of collagen remodeling. GLCM features capture local intensity relationships, wavelet and Fourier analyses quantify structural complexity, tensor metrics emphasize collagen fibers with high intensity and directionality, and Hough-based features characterize local, directional organization. These features do not operate in isolation; rather, their combined behavior provides a multidimensional representation of each tissue state. For example, the coexistence of high coherence fraction with elevated high frequency power in the NR group potentially suggests restoration of global alignment alongside persistent microstructural heterogeneity, consistent with partial but incomplete reversal of fibrosis.

This integrative interpretability is a key strength of the proposed framework. By grounding each feature in a biologically meaningful scope while maintaining tabular compatibility with ML models, the pipeline enables not only accurate classification but also physiological insights into the structural changes underlying disease progression and therapeutic response.

## 4. Discussion

This study presents an interpretable SHG feature-extraction pipeline for quantitative characterization of fibrotic remodeling in murine nonischemic HF and NR. We developed 87 rotationally invariant, biologically interpretable features spanning pixel-level, textural, morphological, and structural domains to quantify collagen density, alignment, and microstructural organization. Using multiple tabular ML models with LOSO cross-validation, the pipeline achieved strong classification performance (AUC: 0.84-0.99), with the highest accuracy obtained using a random forest classifier. These findings demonstrate that normal myocardium, HF, and NR exhibit distinct SHG-derived structural signatures, although further validation is needed before clinical diagnostic or treatment-monitoring applications.

Our feature set substantially outperformed established SHG analysis tools, including CT-FIRE and TWOMBLI, demonstrating the value of capturing multiscale ECM architecture beyond fiber-level descriptors. Feature importance analysis showed that discrimination arose from complementary measures of collagen organization, heterogeneity, and directional coherence, underscoring the advantage of integrating diverse, interpretable features to characterize disease progression and recovery.

Feature filtering was performed before model training to reduce redundancy and mitigate overfitting. Two approaches were evaluated: PCC-based pruning to remove highly collinear features while preserving interpretability, and PCA to project features into lower-dimensional orthogonal space. PCC filtering achieved the best performance with the RF classifier (AUC > 0.99), whereas PCA provided modest improvements for models more sensitive to multicollinearity, such as LDA and DT.

Models were evaluated using LOSO CV at the murine specimen level, ensuring that all FOVs from a given specimen were held out together. This reduces information leakage from tissue architecture and provides a more rigorous assessment than random FOV-level splitting. However, because the study included only 15 independent specimens, these performance estimates remain preliminary and require confirmation through larger cohorts, confidence intervals, specimen-level aggregation, and external validation. Model interpretability was preserved through feature- and group-level importance analyses of the RF classifier.

Our results further demonstrate that SHG-derived features can identify a structural state consistent with NR. Successful discrimination among control, HF, and NR indicates that withdrawal of the HF-inducing stimulus promotes partial, but incomplete, reversal of pathological ECM remodeling. The extracted features consistently captured complementary aspects of collagen organization, indicating a transition from the heterogeneous, aligned architecture of healthy myocardium to a more homogeneous, disorganized, and structurally simplified matrix in HF, with partial restoration following recovery.

Despite the limited dataset (75 FOVs from 15 murine specimens), the proposed feature set produced classification accuracies well above the 33% chance level expected for a balanced three-class task and outperformed two widely used SHG analysis tools, supporting the value of multiscale, interpretable feature extraction. Nevertheless, because the biologically independent sample size is determined by the number of specimens rather than FOVs, these findings should be interpreted cautiously and validated in larger cohorts.

The relative importance of individual features is expected to be tissue specific. In cardiac tissue, the highly organized, anisotropic collagen architecture favors wavelet- and tensor-based metrics that quantify directional organization, complemented by GLCM and Fourier features that capture spatial texture and frequency content. Together, these complementary descriptors provide a robust representation of cardiac ECM remodeling.

Although DL models such as CNNs and vision transformers (ViTs) are considered state of the art in many computer vision medical applications, including various pathology and radiology implementations, there are numerous advantages and limitations compared to tabular models [43]. Perhaps most importantly in the context of in vivo diagnostics using multimodal biophotonics is biological interpretability. While CNN models leverage hundreds, thousands, or even millions of embeddings analogous to our handcrafted features, these embeddings are not readily interpretable to the human eye. Recent developments in DL saliency mapping have attempted to address this issue [44].

Data requirements also differ by orders of magnitude. Structured data classifiers typically require hundreds to thousands of images, whereas DL models like CNNs or ViTs may demand millions of images to avoid overfitting [43]. Although the latter can offer higher performance, it may require thousands of unique patient samples, an unfeasible expectation for most clinical studies. Furthermore, images acquired can come from different systems with varying optical configurations, pixel dimensions, and excitation wavelengths. Therefore, a lightweight model retrainable on minimal data using our curated feature set offers exceptional promise for broad clinical generalizability and equitable distribution of such technologies.

Lastly, a tabular-based model offers multimodal incorporation with immediate interpretability. For example, features derived from patient demographics, clinical presentation or from the use of multimodal optical imaging approaches could be feasibly implemented into the existing framework for increased precision and diagnostic insights. The use of multimodal label-free imaging, specifically, other nonlinear optical microscopy techniques such as two-photon autofluorescence microscopy (TPAF) and coherent anti-Stokes Raman scattering microscopy (CARS) in addition to SHG imaging could provide complimentary information and the inclusion of features derived from these techniques will be a part of our future work [45].

Several features within the pipeline rely on user defined hyperparameters. For example, metrics such as FFT anisotropy, Hough angle distributions, and Hough field descriptors depend on windowing to generate orientation histograms or local structural summaries. The selected window size directly influences the scale of analysis and therefore the biological interpretation of collagen organization. Smaller windows increase spatial resolution and improve quantification of local tissue heterogeneity but may sacrifice sensitivity to longer range structural relationships and global order. Conversely, larger windows better preserve global architectural information while reducing sensitivity to microstructural variability. Future work will incorporate automated hyperparameter optimization to systematically determine window size, wavelet decomposition, GLCM distance, and related parameters based on tissue-specific model performance. Such an approach will improve reproducibility, reduce user-dependent bias, and provide a quantitative basis for parameter selection.

Many features in the current pipeline could be extended to three-dimensional analysis. Benefits of 3D translation include increased spatial resolution with more embedded information. One of the drawbacks with using MIP images in 2D is that some FOVs, as shown in our cardiac example, can be dependent on the tissue cut. For example, if the collagen fibers are sliced axially, they will appear as numerous circles as opposed to fine, parallel lines. 3D feature extraction would in theory be less vulnerable to this discrepancy given the additional dimensionality. The tradeoff would be computing time for the initial feature extraction step; however, this is already quite low for MIP feature extraction, so it would stand to reason that this could be feasibly implemented in clinical settings where features are only being extracted from a handful of FOVs at a time.

For this study, manual segmentation was performed to show proof-of-concept to reduce as many variables as possible. Restricting the anatomical site to adjacent FOVs comprising the LV free wall with longitudinally oriented fibers maximized explainability and consistency across models. Future implementation of texture feature-guided K clustering could be utilized to automatically segment tissue and remove any ambiguity or user input [18].

A key limitation is that the SHG images acquired in this study do not distinguish myosin from collagen in cardiac tissue; therefore, the reported results reflect the combined SHG signal from both components. Accordingly, feature interpretations should be framed as SHG-derived fibrillar structural remodeling unless supported by co-registered collagen stains, biochemical fibrosis assays, or polarization-sensitive SHG. Prior studies have demonstrated that polarization-sensitive SHG can help distinguish myosin and collagen, and incorporation of this approach will be pursued in future work.

## Acknowledgment

The authors would like to thank Ms. Rucha Dave for her technical assistance.

## Data availability

Data underlying the results presented in this paper are not publicly available at this time but may be obtained from the authors upon reasonable request. Given the absence of established standards for these measurements, public release of the data and code would not offer additional interpretive value for this work

