## Supplementary information for "Interpretable computational SHG microscopy enables quantitative tissue phenotyping of cardiac remodeling in nonischemic heart failure"

### INTERPRETABLE SHG MICROSCOPY FEATURE-EXTRACTION FRAMEWORK FOR QUANTITATIVE ASSESSMENT OF CARDIAC REMODELING IN NONISCHEMIC HEART FAILURE: SUPPLEMENTAL DOCUMENT

**Table S1. Summary of all 87 features derived from different analyses.**

| Domain | Analysis | Feature Type | Method(s) | Individual Features |
|---|---|---|---|---|
| Spatial | Intensity | First-order statistics | Raw intensity | mean_int, std_int, skew_int, kurt_int, entropy_int |
| | SHG-derived | Signal intensity | Collagen area fraction | CAF |
| | GLCM | Co-occurrence texture | Autocorrelation, Contrast, Dissimilarity, Homogeneity, Max Probability, Energy, Entropy | autocorr, contrast, dissimil, homogeneity, maxprob, energy_GLCM, entropy_GLCM |
| | | Cluster-based features | Cluster prominence, shade, and tendency | clusterp, clusters, clustert |
| | | Statistical measures | Correlation, sum of squares, inverse variance | correlat, sumsqvar, inv_var |
| | | Sum and difference matrices | Sum distribution, difference distribution | sumavg, sument, diffent, diffvar |
| | | Information measures of correlation (IMC) | IMC-1, IMC-2 | inf1h, inf2h |
| | | Inverse difference variants | Inverse difference normalized (INN), inverse difference moment normalized (IDMN) | invdn, invdmn |
| | OD-GLCM | Angular GLCM | Cumulative correlation | Std_cumcorr |
| | GLRLM | Run length matrix | Short vs. long run emphasis, high grayscale intensity emphasis | LRE, GLN, RLN, RP, HGRE, LRHGE |
| | Tensor | Tensor-based orientation | Anisotropy-weighted intensity (AWI) | mean_awi, std_awi, skew_awi, kurt_awi, entropy_awi |

| | | | | |
|---|---|---|---|---|
| | | | Degree of isotropy (DOI) | mean_DOI, std_DOI, skew_DOI, kurt_DOI, entropy_DOI |
| | | | Preferred orientation (PO) | std_PO, skew_PO, kurt_PO, entropy_PO |
| | | Tensor magnitude | Gradient energy (trace) | mean_trace, std_trace, skew_trace, kurt_trace, entropy_trace |
| | Fractal | Fractal geometry | Fractal dimension (binary and grayscale) | fd_bin, fd_gray |
| | | Fractal texture | Lacunarity (binary and grayscale) | lac_bin, lac_gray |
| Frequency | FFT | Power spectrum | Total power | total_power |
| | | | Low/mid/high frequency power | low_freq_power, mid_freq_power, high_freq_power |
| | | | Weighted frequency center | WFC |
| | | Directional texture | Radial summation (anisotropy) | anisotropy |
| Spatial & Frequency | Wavelet | Wavelet coarse energy | Total energy | E_total |
| | | | Approximation coefficients | meanA, E_A |
| | | Wavelet detail | Detail energies | L1_total, L2_total |
| | | Wavelet energy balance | Energy ratios | Ratio_L1L2, ratio_LL_to_details |
| | | Wavelet texture | Wavelet entropy | WavEntropy, entA, ent_L1, ent_L2 |
| Hough Domain | Hough Transform | Line detection | Hough standard deviation | hough_std |
| | Hough field | Orientation reliability | Coherence fraction | coherence_fraction |
| | | Orientation dispersion | Orientation dispersion (degrees) | ori_dispersion_deg |
| | | Local axial order | Local alignment magnitude | mean_Slocal, std_Slocal, skew_Slocal, |

| | | | |
|---|---|---|---|
| | | | kurt_Slocal, ent_Slocal |
| | Neighborhood alignment | Neighborhood alignment coherence | mean_nalign, std_nalign, skew_nalign, kurt_nalign, ent_nalign |

**Table S2. Summary of CT-FIRE feature set.**

| Feature | Brief mathematical definition (fiber-level) | FOV-level aggregation (what is stored) | Biological meaning (interpretation) |
|---|---|---|---|
| ctf_fiber_num | N = number of extracted fibers (count) | Scalar N per FOV | Fiber abundance / density proxy (more fibers detected = more collagen structures in FOV) |
| ctf_len_mean | Mean of length histogram | mean(L_i) | Average fiber extent (longer fibers may indicate more continuous collagen bundles) |
| ctf_len_std | Standard deviation of length histogram | std(L_i) | Heterogeneity in fiber lengths (mixed short/long fibers) |
| ctf_len_median | Median of length histogram | median(L_i) | Typical fiber length |
| ctf_len_iqr | Interquartile range of length histogram | IQR(L_i) | Robust spread of fiber lengths (length variability) |
| ctf_len_min | Minimum of length histogram | min(L_i) | Shortest detected fibers (fragmentation / detection of small segments) |
| ctf_len_max | Maximum of length histogram | max(L_i) | Longest detected fibers (presence of long continuous fibers) |
| ctf_len_cv | Ratio of std to mean | CV = std(L_i)/mean(L_i) | Relative length variability (scale-normalized heterogeneity) |
| ctf_len_total | Sum of length histogram | Total = Σ L_i (or provided totL) | Total collagen fiber content proxy within FOV |

| | | | |
|---|---|---|---|
| ctf_angle_mean | Mean of angle histogram | mean(θ_i) | Dominant orientation direction (preferred alignment axis) |
| ctf_angle_std | Standard deviation of angle histogram | std(θ_i) | Dispersion of orientations (higher = less aligned / more disordered) |
| ctf_angle_nematic_S | S = \|⟨cos(2θ)⟩\| | Scalar S per FOV | Degree of alignment (0 = isotropic, 1 = highly aligned collagen) |
| ctf_xlink_count | X = number of detected crosslinks | Scalar X per FOV | Crosslinking / branching events proxy (network connectivity) |
| ctf_xlink_density | X as above; A = area/volume proxy | Density = X/A (or provided xlinkdens) | Connectivity normalized to size; crosslink concentration proxy |
| ctf_xlinkspace_mean | Mean of crosslink histogram | mean(s_j) | Typical spacing between crosslinks (network mesh size proxy) |
| ctf_xlinkspace_std | Standard deviation of crosslink histogram | std(s_j) | Variability of mesh size / crosslink spacing |
| ctf_radius_mean | Mean of proxy fiber radius histogram | mean(r_k) | Average thickness proxy (thicker fibers often reflect higher collagen bundling) |
| ctf_radius_std | Standard deviation of proxy fiber radius histogram | std(r_k) | Thickness heterogeneity across sampled locations |
| ctf_radius_median | Median of proxy radius histogram | median(r_k) | Typical thickness |
| ctf_straight_mean | Mean of straightness histogram | mean(straightness_i) | Average waviness/straightness (lower = more crimped or tortuous fibers) |
| ctf_straight_std | Standard deviation of straightness histogram | std(straightness_i) | Heterogeneity in waviness across fibers |
| ctf_fiber_radius_mean | Mean of individual fiber radius histogram | mean(r̄_i) | Average per-fiber thickness (fiber-level thickness proxy) |
| ctf_fiber_radius_std | Standard deviation of individual | std(r̄_i) | Between-fiber thickness heterogeneity (mix of thin vs thick fibers) |

| | |
|---|---|
| | fiber radius histogram |

**Table S3. Summary of median values for control, heart failure, and treatment murine models with associated p values.**

| Feature | Control | Heart Failure | Treatment | Control vs heart failure | Control vs treatment | Heart failure vs treatment |
|---|---|---|---|---|---|---|
| Invdn | 0.959 | 0.971 | 0.929 | **** | **** | **** |
| L1 total | 355.3 | 255.1 | 1596.0 | **** | **** | **** |
| Mean AWI | 0.138 | 0.059 | 0.072 | **** | **** | * |
| High frequency power | 9.91E6 | 5.10E6 | 6.18E7 | **** | **** | **** |
| Coherence fraction | 0.720 | 0.342 | 0.751 | **** | *** | **** |

*p < 0.05, **p < 0.01, ***p < 0.001, **** p < 0.0001

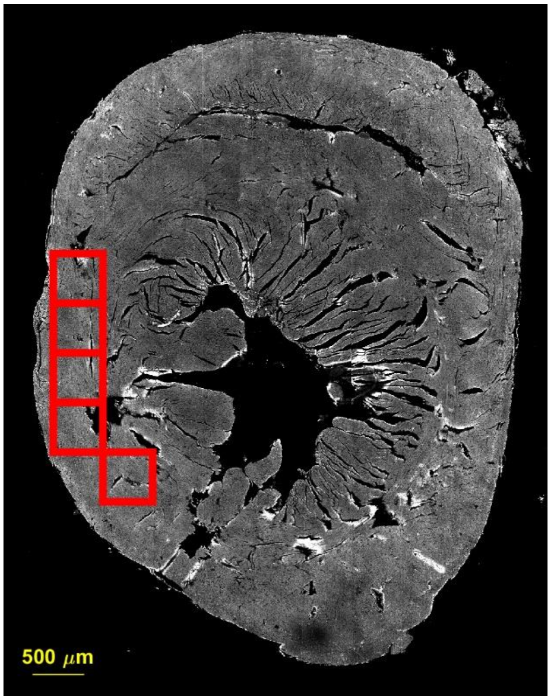


Fig. S1. Control sample 1 segmentations.

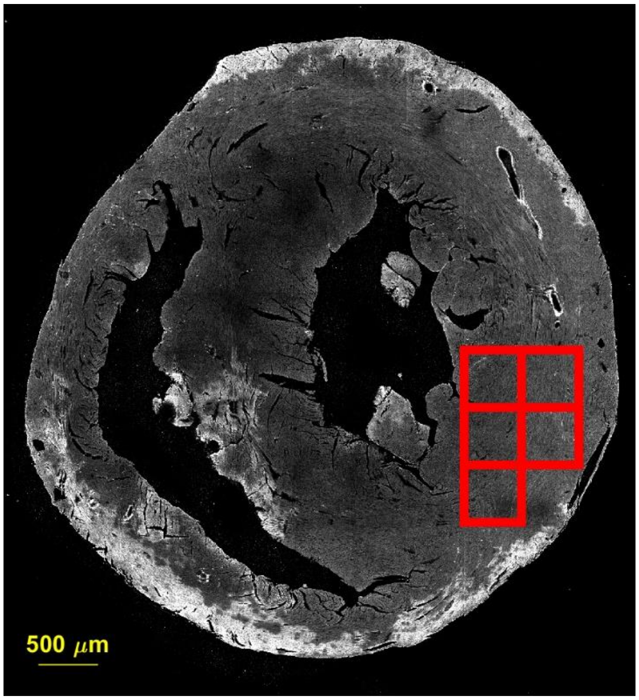


Fig. 2. Control sample 2 segmentations.

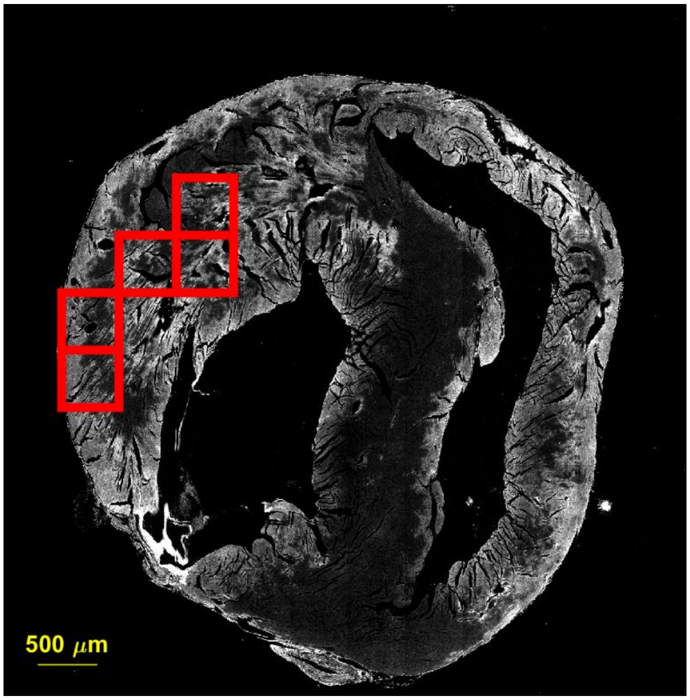


Fig. S3. Control sample 3 segmentations.

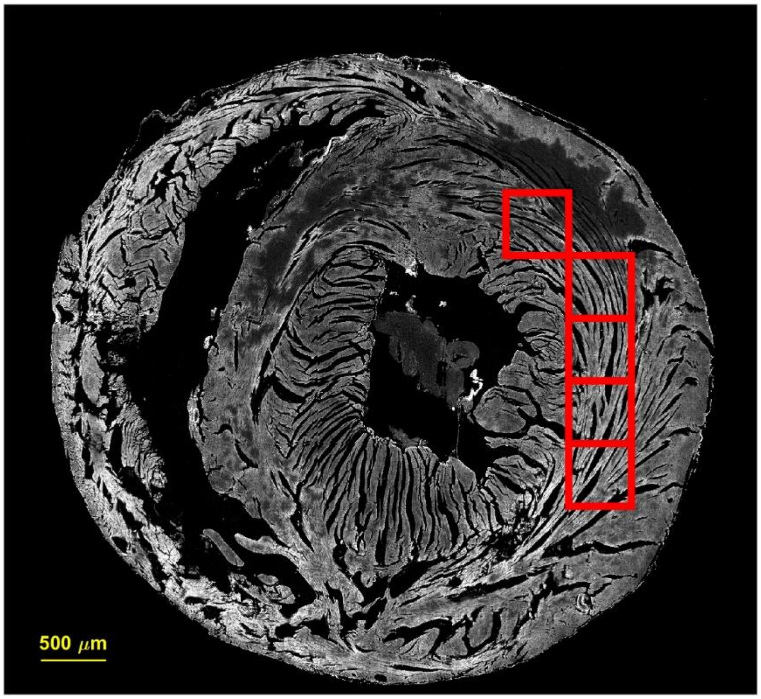


Fig. S4. Control sample 4 segmentations.

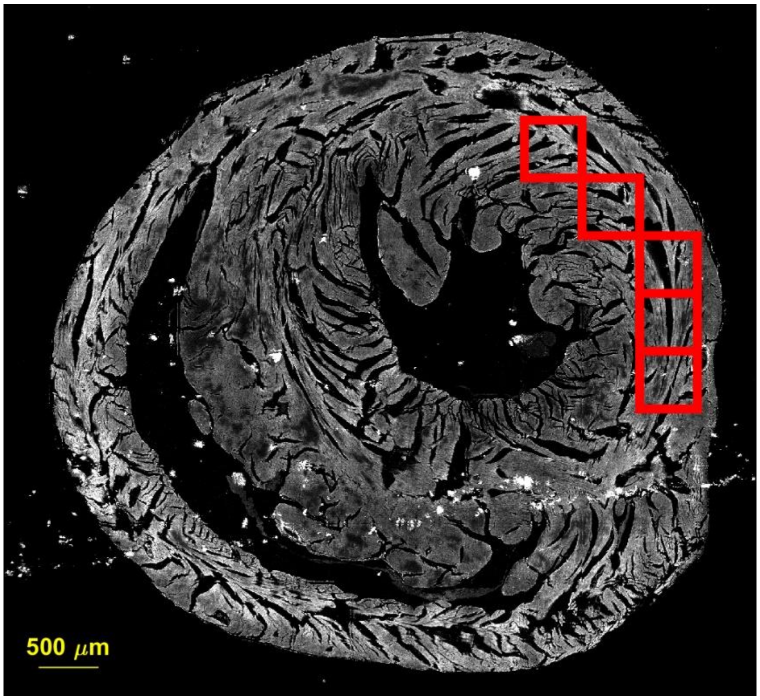


Fig. S5. Control sample 5 segmentations..

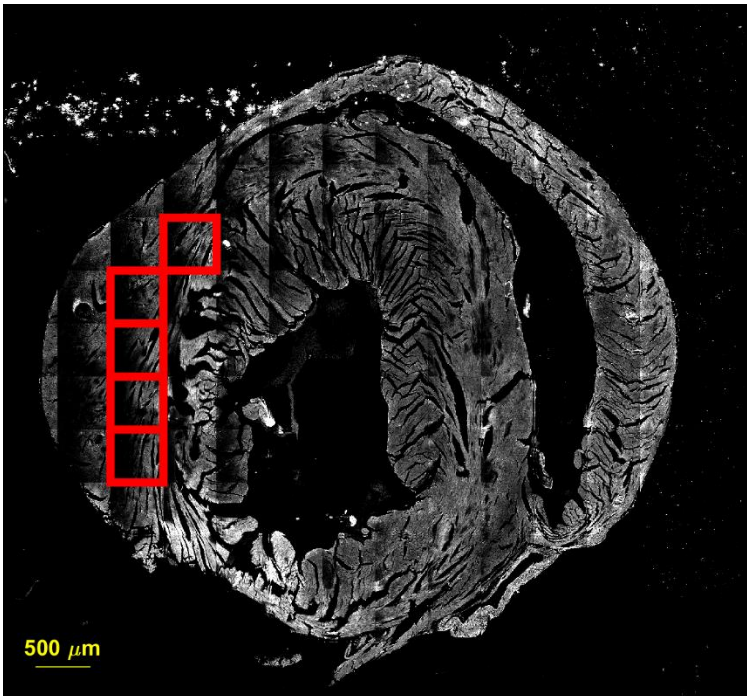


Fig. S6. Heart failure sample 1 segmentations.

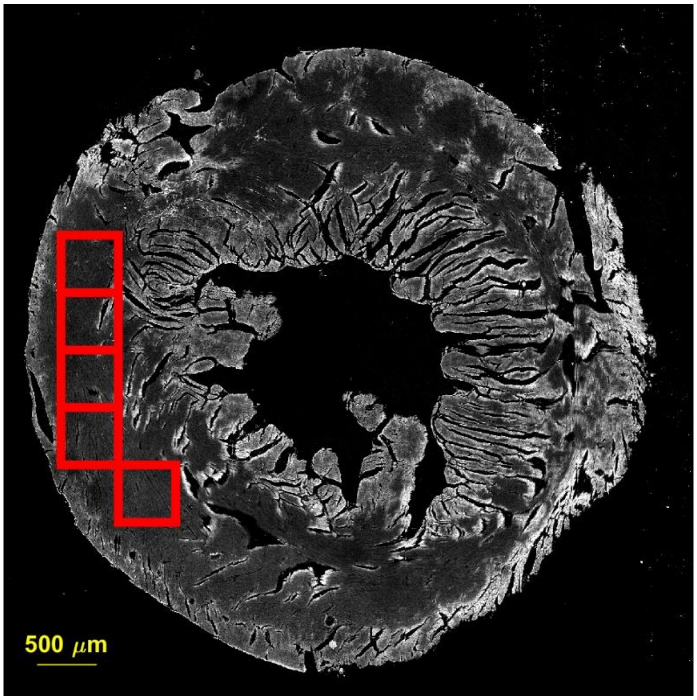


Fig. S7. Heart failure sample 2 segmentations.

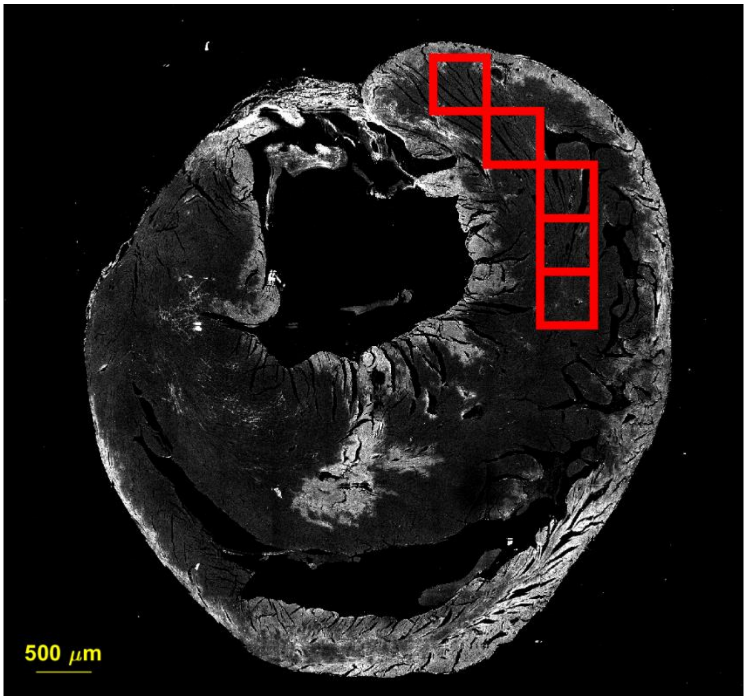


Fig. S8. Heart failure sample 3 segmentations.

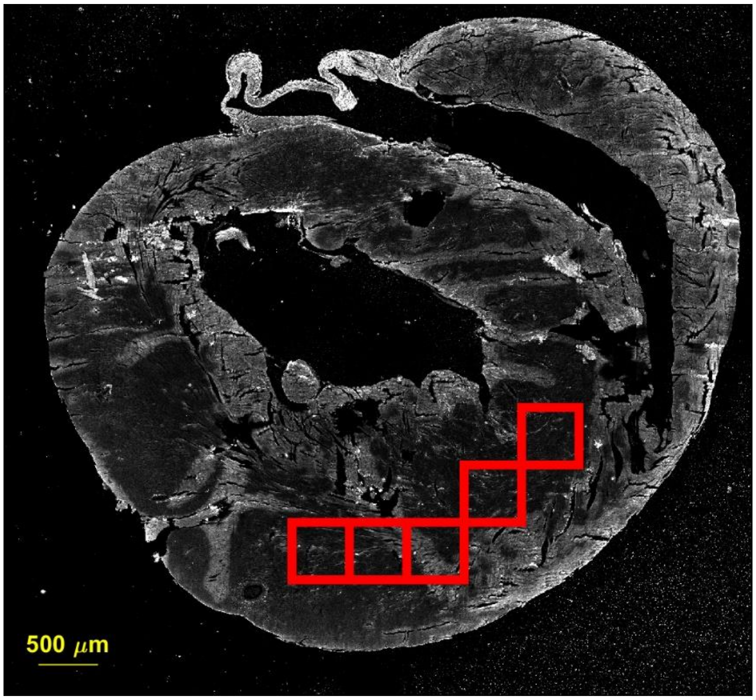


Fig. S9. Heart failure sample 4 segmentations.

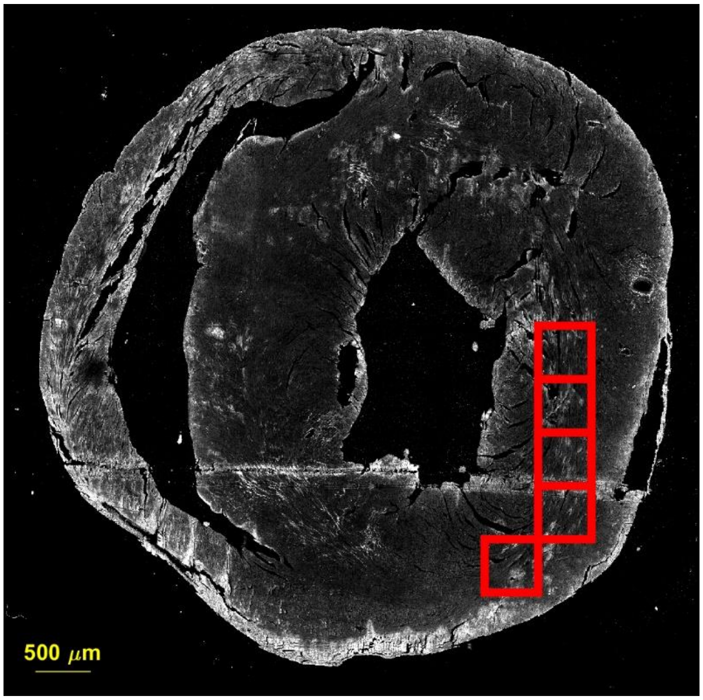


Fig. S10. Heart failure sample 5 segmentations.

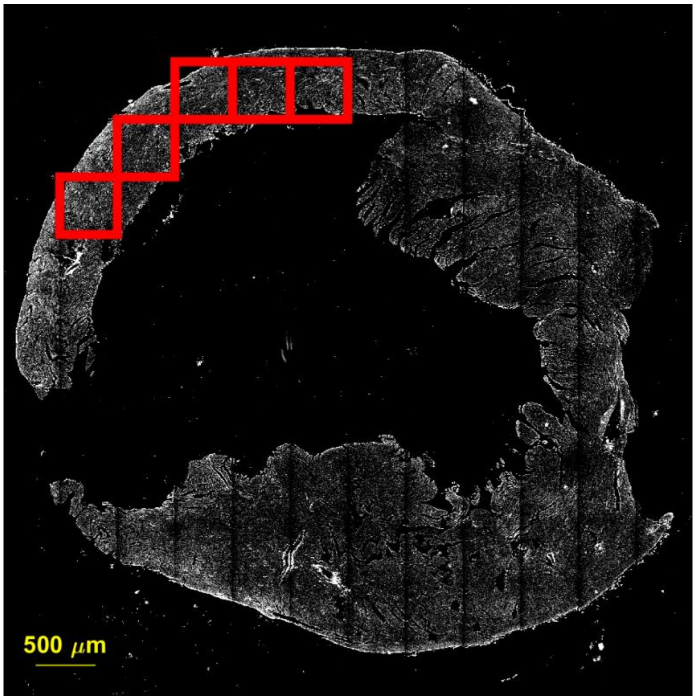


Fig. S11. Natural recovery sample 1 segmentations.

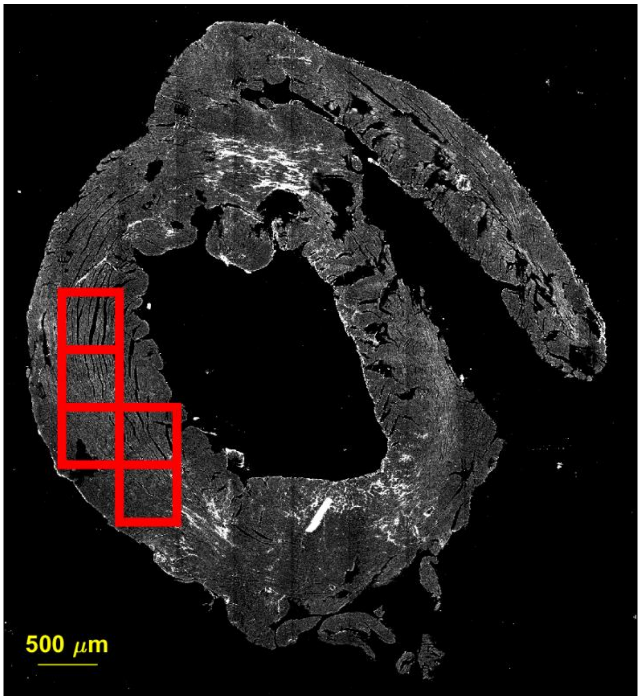


Fig. S12. Natural recovery sample 2 segmentations.

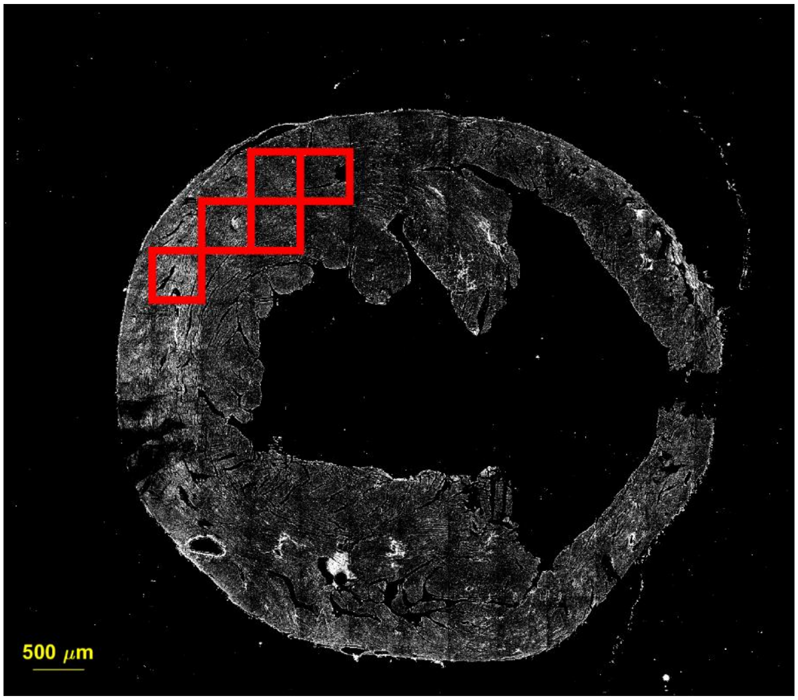


Fig. S13. Natural recovery sample 3 segmentations.

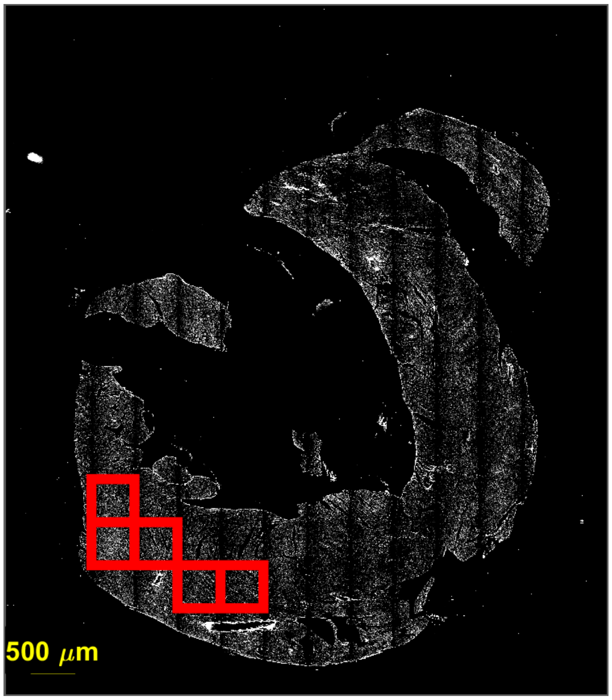


Fig. S14. Natural recovery sample 4 segmentations.

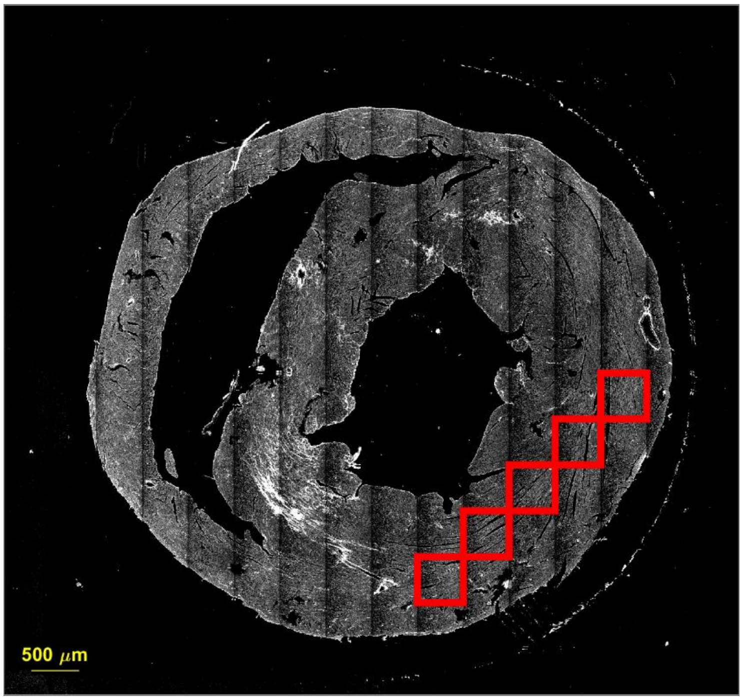


Fig. S15. Natural recovery sample 5 segmentations.

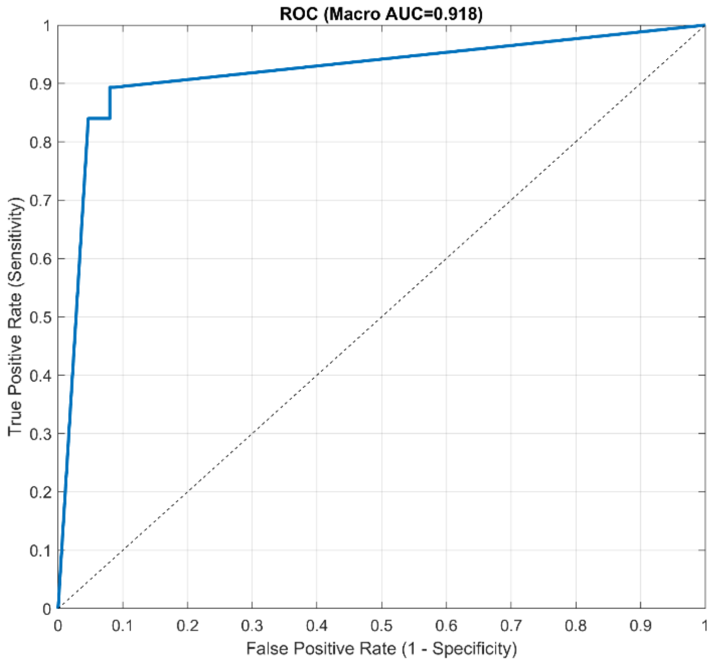

Fig. S16. ROC-AUC plot for our features using PCA filtering and DT model.

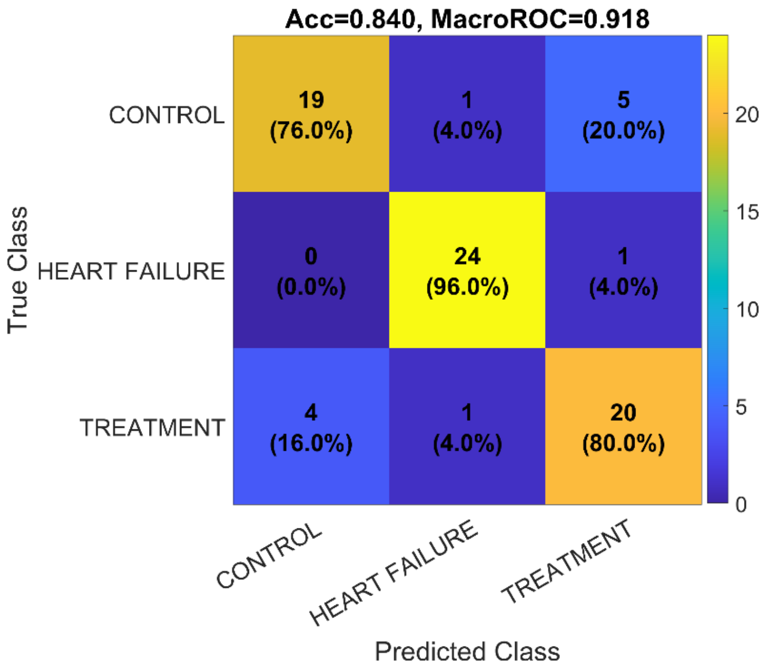

Fig. S17. Confusion matrix for our features using PCA filtering and DT model.

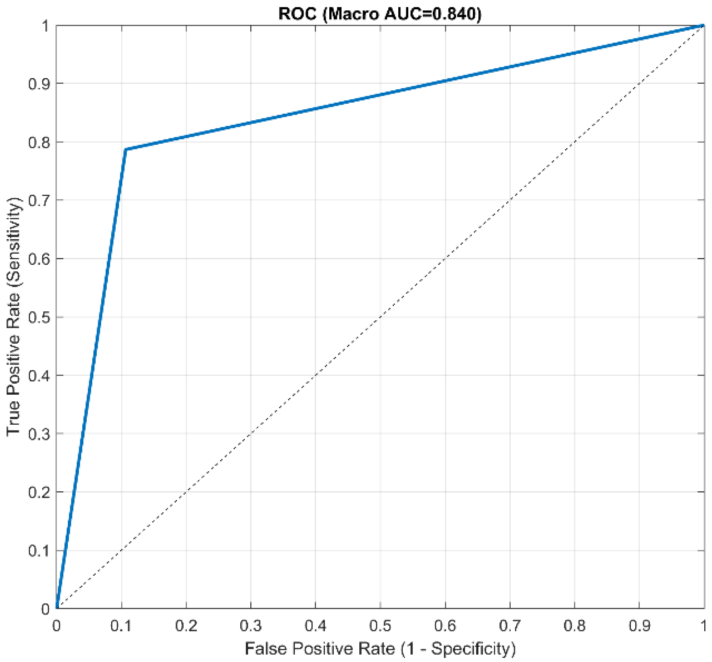


Fig. S18. ROC-AUC plot for our features using PCC filtering and DT model.

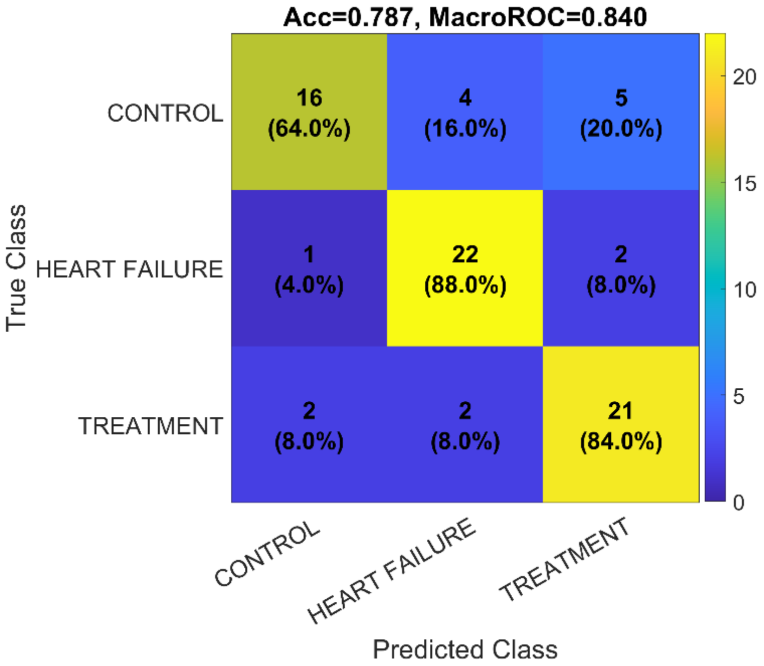


Fig. S19. Confusion matrix for our features using PCC filtering and DT model.

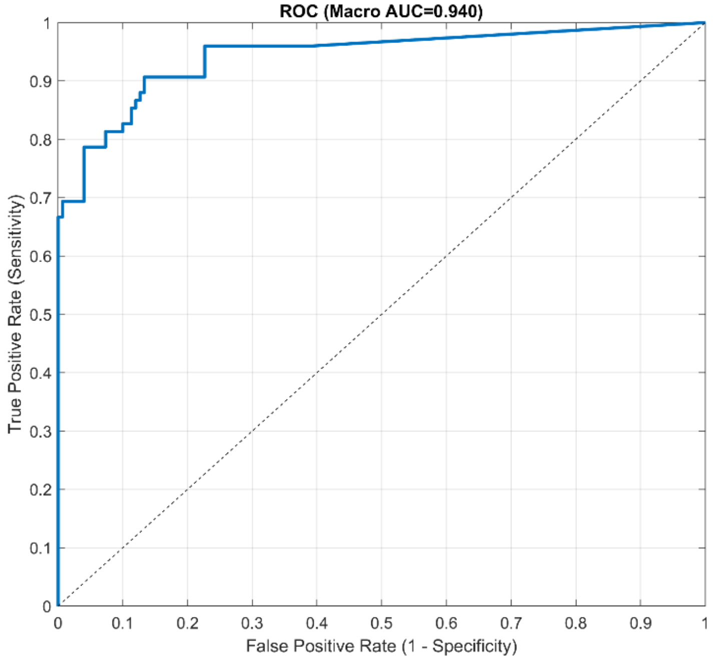


Fig. S20. ROC-AUC plot for our features using PCA filtering and KNN model.

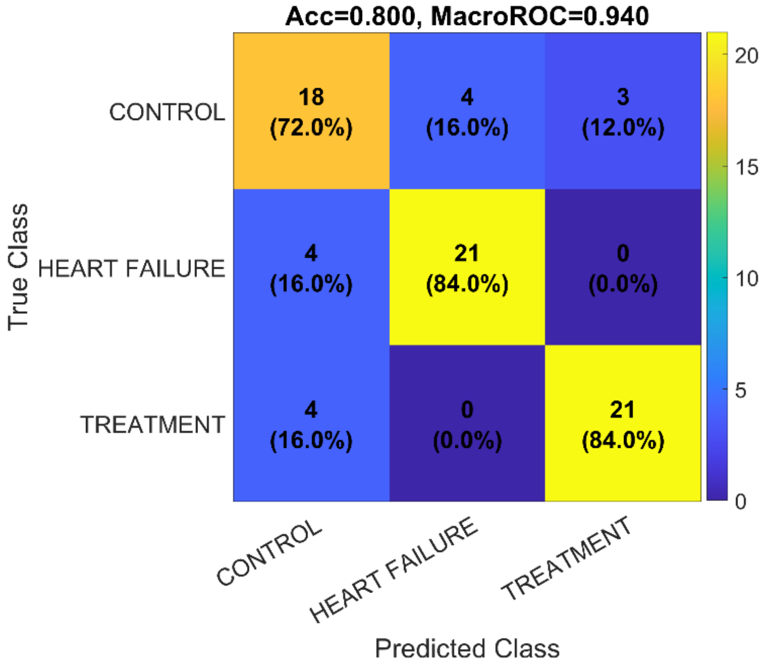


Fig. S21. Confusion matrix for our features using PCA filtering and KNN model.

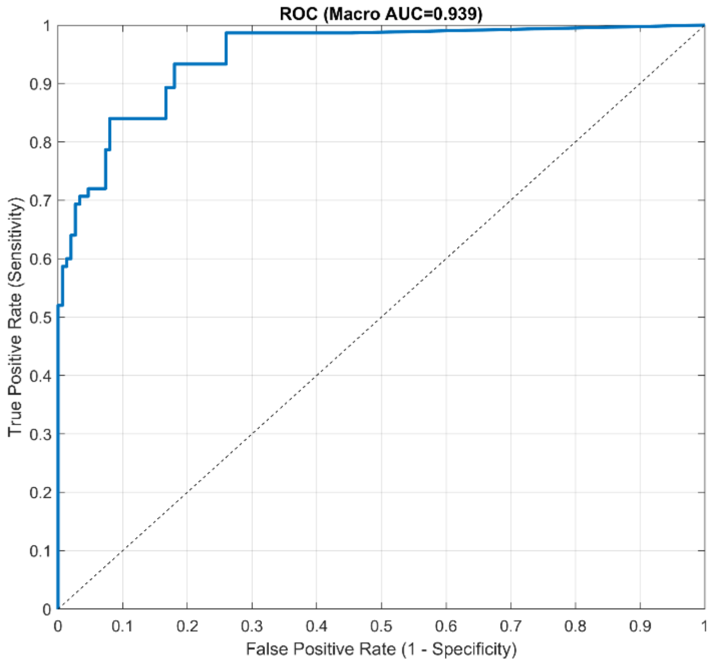

Fig. S22. ROC-AUC plot for our features using PCC filtering and KNN model.

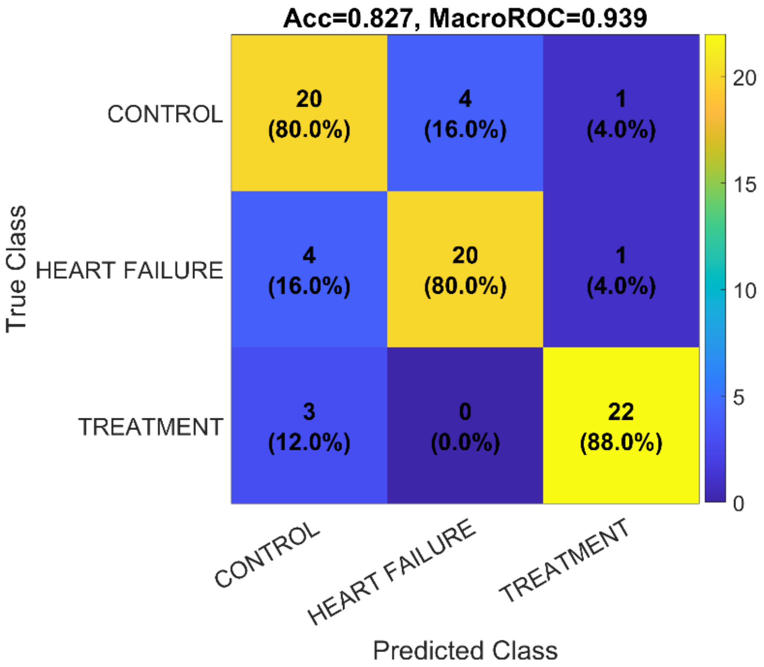

Fig. S23. Confusion matrix for our features using PCC filtering and KNN model.

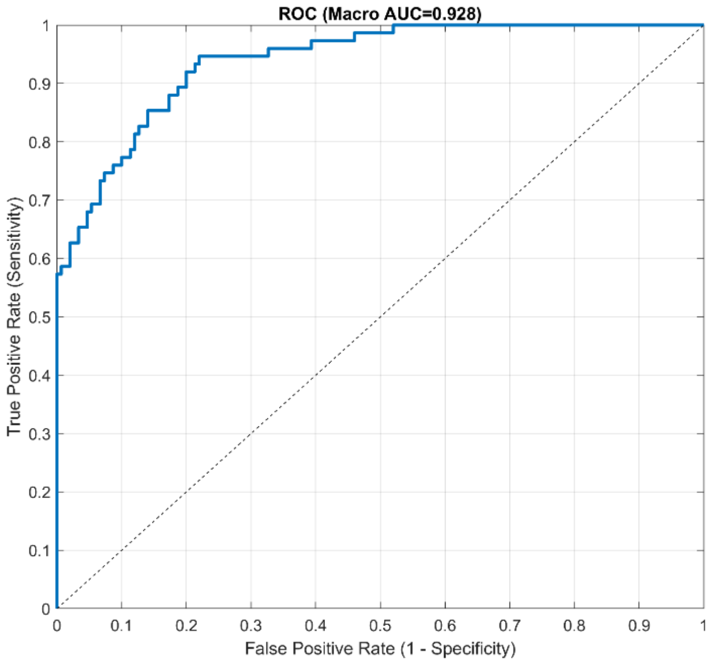


Fig. S24. ROC-AUC plot for our features using PCA filtering and LDA model.

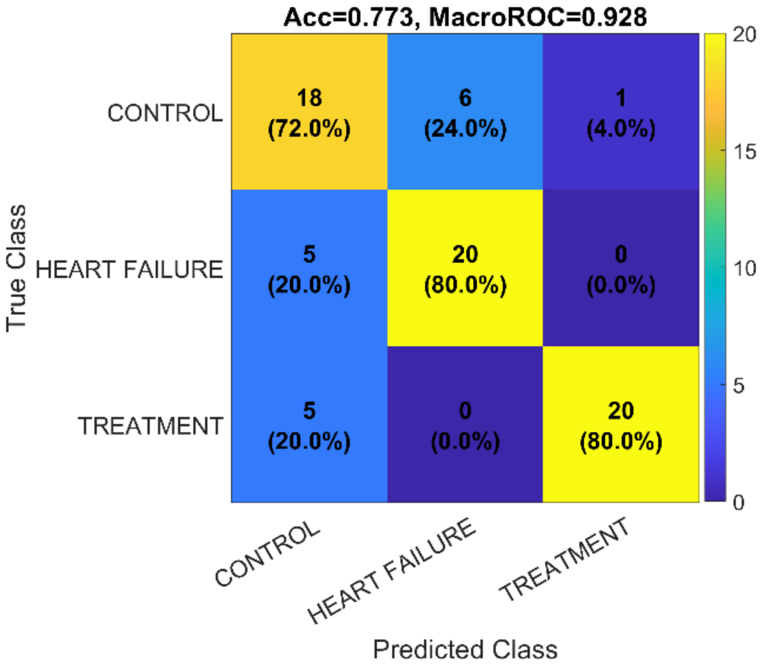


Fig. S25. Confusion matrix for our features using PCA filtering and LDA model.

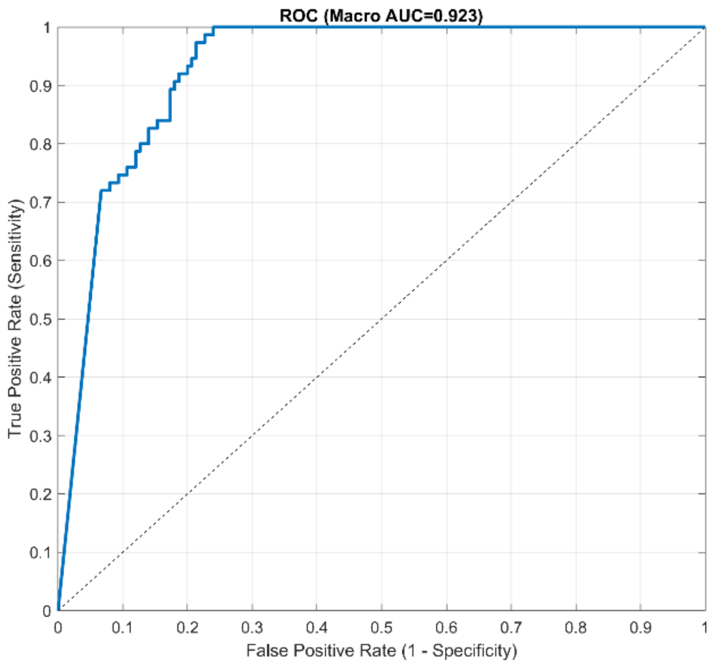


Fig. S26. ROC-AUC plot for our features using PCC filtering and LDA model.

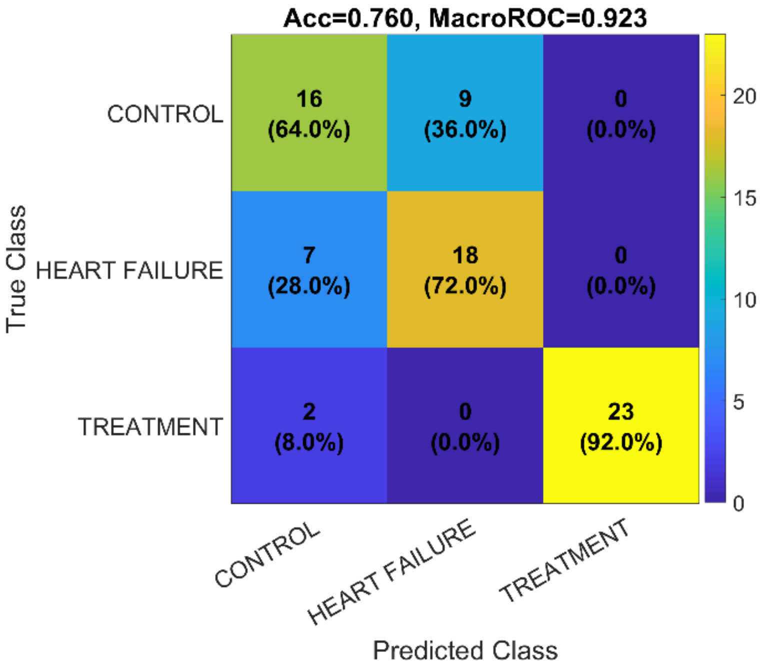


Fig. S27. Confusion matrix for our features using PCC filtering and LDA model.

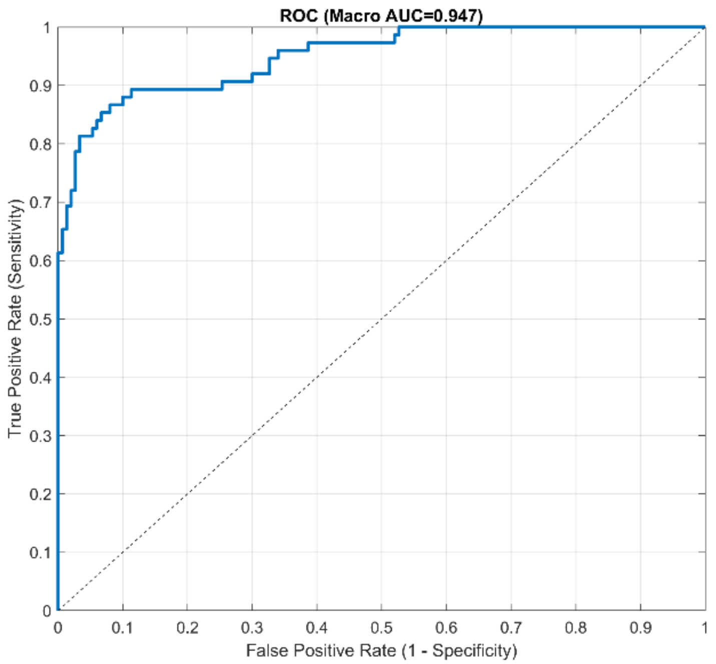


Fig. S28. ROC-AUC plot for our features using PCA filtering and RF model.

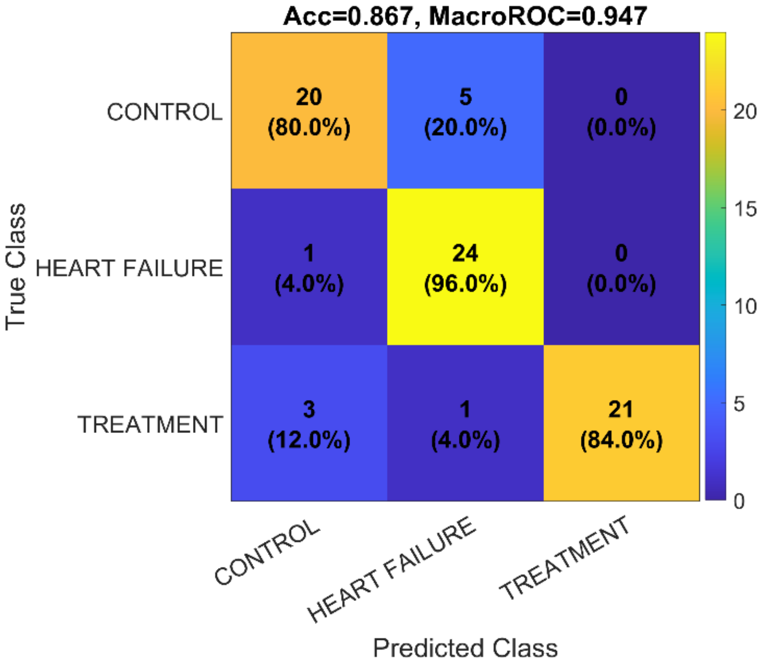


Fig. S29. Confusion matrix for our features using PCA filtering and RF model.

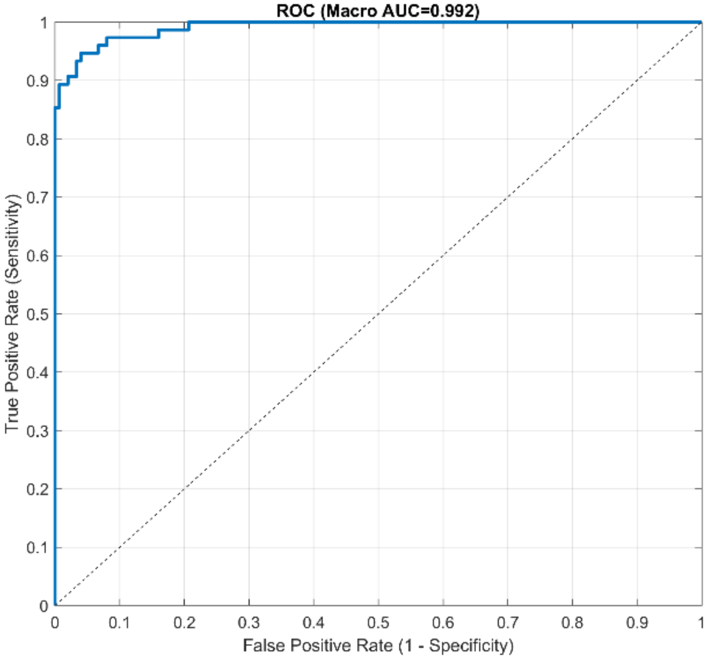


Fig. S30. ROC-AUC plot for our features using PCC filtering and RF model.

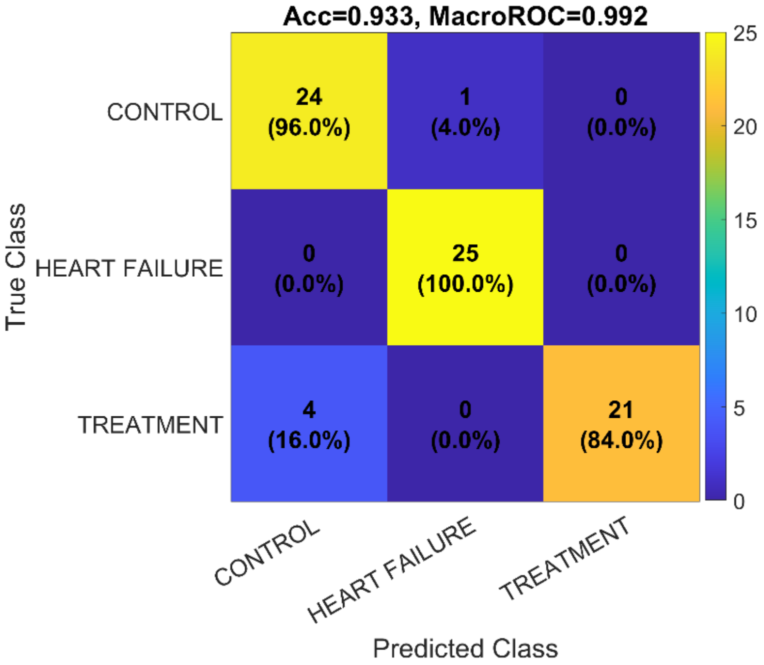


Fig. S31. Confusion matrix for our features using PCC filtering and RF model.

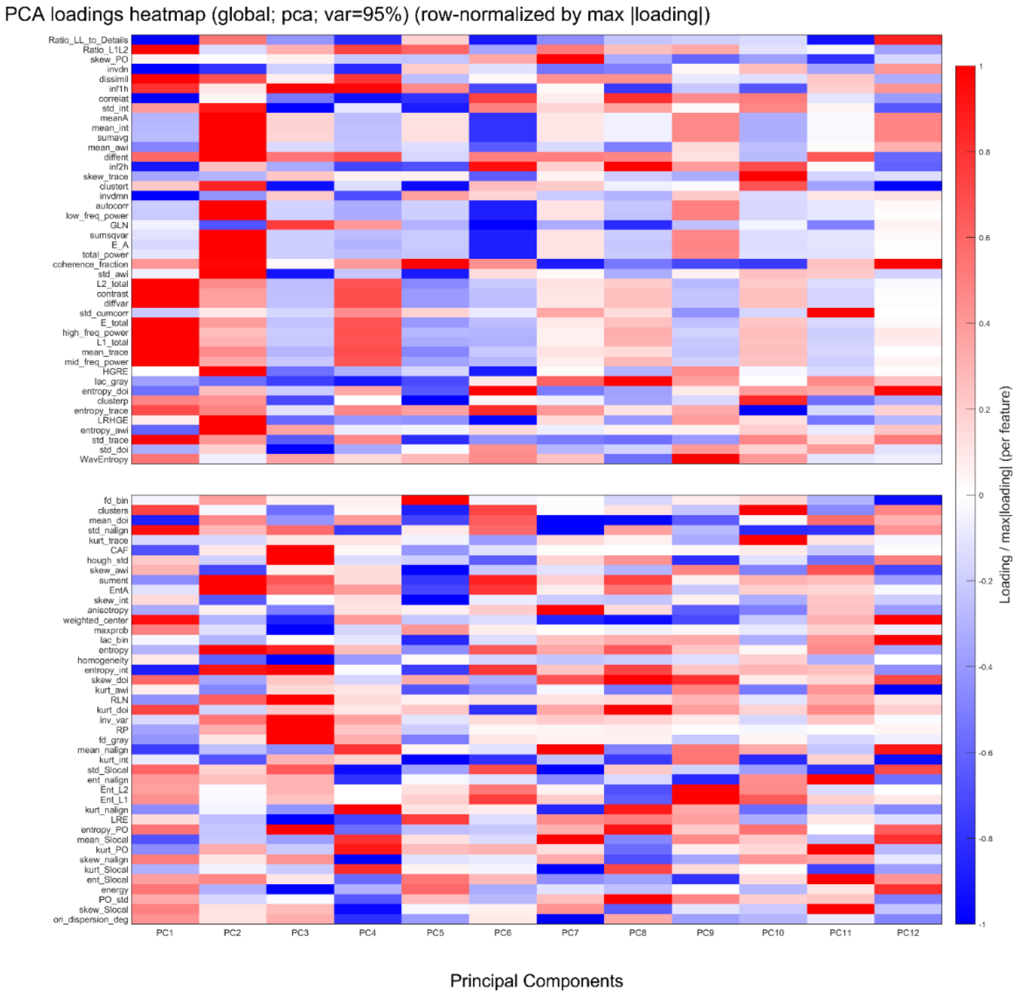


Fig. S32. PCA loadings for RF model.
